# Emergent selectivity for scenes, object properties, and contour statistics in feedforward models of scene-preferring cortex

**DOI:** 10.1101/2021.09.24.461733

**Authors:** Donald Shi Pui Li, Michael F. Bonner

**Affiliations:** Department of Cognitive Science, Johns Hopkins University, Baltimore, MD, USA

**Keywords:** category-selective cortex, visual features, convolutional neural networks, encoding models, fMRI

## Abstract

The scene-preferring portion of the human ventral visual stream, known as the parahippocampal place area (PPA), responds to scenes and landmark objects, which tend to be large in real-world size, fixed in location, and inanimate. However, the PPA also exhibits preferences for low-level contour statistics, including rectilinearity and cardinal orientations, that are not directly predicted by theories of scene- and landmark-selectivity. It is unknown whether these divergent findings of both low- and high-level selectivity in the PPA can be explained by a unified computational theory. To address this issue, we fit feedforward computational models of visual feature coding to the image-evoked fMRI responses of the PPA, and we performed a series of high-throughput experiments on these models. Our findings show that feedforward models of the PPA exhibit emergent selectivity across multiple levels of complexity, giving rise to seemingly high-level preferences for scenes and for objects that are large, spatially fixed, and inanimate/manmade while simultaneously yielding low-level preferences for rectilinear shapes and cardinal orientations. These results reconcile disparate theories of PPA function in a unified model of feedforward feature coding, and they demonstrate how multifaceted selectivity profiles naturally emerge from the feedforward computations of visual cortex and the natural statistics of images.

**SIGNIFICANCE STATEMENT:** Visual neuroscientists characterize cortical selectivity by identifying stimuli that drive regional responses. A perplexing finding is that many higher-order visual regions exhibit selectivity profiles spanning multiple levels of complexity: they respond to highly complex categories, such as scenes and landmarks, but also to surprisingly simplistic features, such as specific contour orientations. Using large-scale computational analyses and human brain imaging, we show how multifaceted selectivity in scene-preferring cortex can emerge from the feedforward, hierarchical coding of visual features. Our work reconciles seemingly divergent findings of selectivity in scene-preferring cortex and suggests that surprisingly simple feedforward feature representations may be central to the category-selective organization of the human visual system.

## INTRODUCTION

A central goal of visual neuroscience is to identify the stimulus properties that selectively drive the responses of neural populations in visual cortex. In high-order visual areas, responses often exhibit complex and unintuitive patterns of multifaceted selectivity for stimulus properties spanning from low-level image features to high-level conceptual attributes^1–4^. Several lines of research suggest that the multifaceted selectivity profiles of higher-order visual regions cannot be reduced to a single level of explanation: neither low-level perceptual factors nor high-level conceptual factors fully account for cortical response preferences^5–9^. Thus, a major challenge for visual neuroscience is to explain how such multifaceted selectivity profiles emerge from the information-processing mechanisms of visual cortex.

The parahippocampal place area (PPA) is a clear example of such multifaceted selectivity. The PPA is a scene-preferring area of the ventral visual stream that responds strongly to spatial scenes^10^. A longstanding hypothesis of the PPA is that it selectively processes large scene elements, including spatial structures and objects, that can serve as navigational landmarks and visual cues for scene categorization^11–17^. This hypothesis is primarily motivated by the strong global signal modulation of the PPA in response to scenes and to objects that are large in real-world size, spatially fixed, and inanimate^13, 16^. However, other work has shown that the responses of the PPA are also modulated by low-level visual features, with a specific preference for high spatial-frequency contours that form rectilinear junctions and are oriented along the cardinal axes^18–20^. Low-level features can even drive the responses of the PPA when presented in minimal stimuli, such as basic geometric shapes, that do not resemble natural scenes or landmarks^19, 20^.

The existence of low-level feature preferences in the PPA has been argued to be inconsistent with theories of landmark- and scene-specialization, and it has sparked a debate over the appropriate level of interpretation for PPA selectivity^2, 6, 17–20^. However, several findings suggest that the response preferences of the PPA cannot be fully explained by low-level features alone. First, the PPA shows a preference for scenes even when they are matched to comparison stimuli on low-level properties^18, 21^. Second, the PPA exhibits a preference for spatial scenes and large objects even in the absence of visual stimulation, when sighted subjects haptically explore miniature scenes or when blind subjects are cued to think of large objects^22, 23^. Nonetheless, the low-level feature preferences of the PPA remain to be explained. Understanding how these low-level preferences square with findings of scene- and landmark-selectivity is critical for developing a complete theory of the PPA, and it may have broader implications for understanding the complex tuning functions of category-selective visual cortex in general.

We explored the possibility that the multifaceted selectivity profile of the PPA can be understood as an emergent property of feedforward, hierarchical computations and the natural statistics of scenes^6, 24^. Our hypothesis was that a small set of feedforward computations is sufficient to produce the high-level selectivity patterns of the PPA and that a direct consequence of these feedforward computations is the emergence of low-level biases for cardinal orientations and rectilinear shapes. Although previous work has speculated that the low-level biases of scene areas arise from selectivity for the perceptual features that are most associated with scenes and landmarks, no studies have directly tested this hypothesis^2, 6^. Furthermore, it has been alternatively speculated that apparent low-level feature preferences in scene areas may instead reflect a tendency for subjects to interpret simplistic rectilinear stimuli as scene-like, which would mean that low-level stimuli only drive scene regions insofar as they are *interpreted* as high-level scenes^18^. It thus remains unknown how to reconcile the disparate selectivity patterns of scene-selective cortex.

A powerful approach for addressing such questions is the use of high-throughput experiments on neural network models of the visual system^24–27^. It has previously been shown that image-computable neural networks can yield remarkably accurate encoding models of category-selective visual cortex, including the PPA^24, 25^, but no studies have directly examined whether selectivity for both high-level scene and object properties and low-level contour statistics can emerge from a unified representational model. To address this question, we fit feedforward neural network models to the scene-evoked fMRI responses of the PPA. We then ran a series of high-throughput, *in-silico* experiments on these models to characterize their selectivity to multiple properties of both natural images and simple geometric stimuli. We found that the feedforward coding of visual features is sufficient to predict fMRI responses in the PPA to natural scenes and objects and to reproduce the selectivity profile of the PPA across multiple levels of complexity, including selectivity for scenes, selectivity for objects that are large, inanimate, manmade, and spatially fixed, and selectivity for the low-level contour statistics of rectilinearity and cardinal orientations. These findings suggest that the multifaceted selectivity profile of the PPA may naturally emerge from the feedforward coding of visual features and the statistical regularities of images.

## RESULTS

### Encoding model of mid-level feature tuning

We used convolutional neural networks (CNNs) and fMRI data to create image-computable voxelwise encoding models of mid-level feature tuning in the PPA. CNNs are theoretical models of the core information-processing mechanisms implemented by biological neural populations, and they are the leading computational models of human visual cortex^28^, including scene-selective areas^24, 25, 29^. They perform a set of biologically plausible mathematical operations, and their hierarchical, convolutional architecture is inspired by the primate visual system. CNNs take images as inputs and pass them through a hierarchy of nonlinear transformations whose final outputs support image classification (after model training). A major strength of CNNs is that they make explicit predictions about the stimulus transformations that may occur along the processing hierarchy of visual cortex. This makes CNNs ideally suited for testing theories about the computational basis of multifaceted selectivity.

We developed voxelwise computational models of visual feature tuning by mapping the outputs of a pre-trained feedforward CNN to fMRI responses in four subjects from the BOLD5000 dataset who viewed between 2,952 and 4,916 unique natural scene images depicting real-world environments and objects^30^. We used the AlexNet CNN architecture pre-trained on scene images (Places365)^31^. The first five layers of AlexNet are convolutional layers, whose units receive inputs from spatially local regions of the previous layer, like the spatial receptive field structure of visual cortex. Each unit performs a linear-nonlinear operation in which it computes a weighted linear sum of its inputs followed by a nonlinear activation function (specifically, a rectified linear threshold). The weights on the inputs for each unit define a type of feature channel, and each convolutional layer contains a set of feature channels that are replicated with the same set of weights over the entire image. Our modelling procedure involved pooling and reweighting of the CNN responses from a convolutional layer to predict the image-evoked fMRI responses in the BOLD5000 dataset (Fig. 1A). We were specifically interested in characterizing feature tuning in the PPA rather than retinotopic biases. We therefore created a set of fully spatially invariant feature activations by applying global max pooling across all spatial locations for each feature channel in the layer. The outputs of this global max pooling operation were passed to a linear regression layer that we trained to predict fMRI responses as a weighted sum of feature activations. We used regularized regression to develop sparse models of feature tuning that focus on the most informative features for each voxel. We compared cross-validated performance when using LASSO (L1 penalized), which encourages sparse models, ridge (L2 penalized), and ordinary least squares (OLS) regression. We found that LASSO outperformed both ridge and OLS regression, suggesting that our models of feature tuning benefited from the inclusion of a regularization term that pushes a portion of the regression weights to zero (Supplementary Fig. 1).

**Figure 1.**
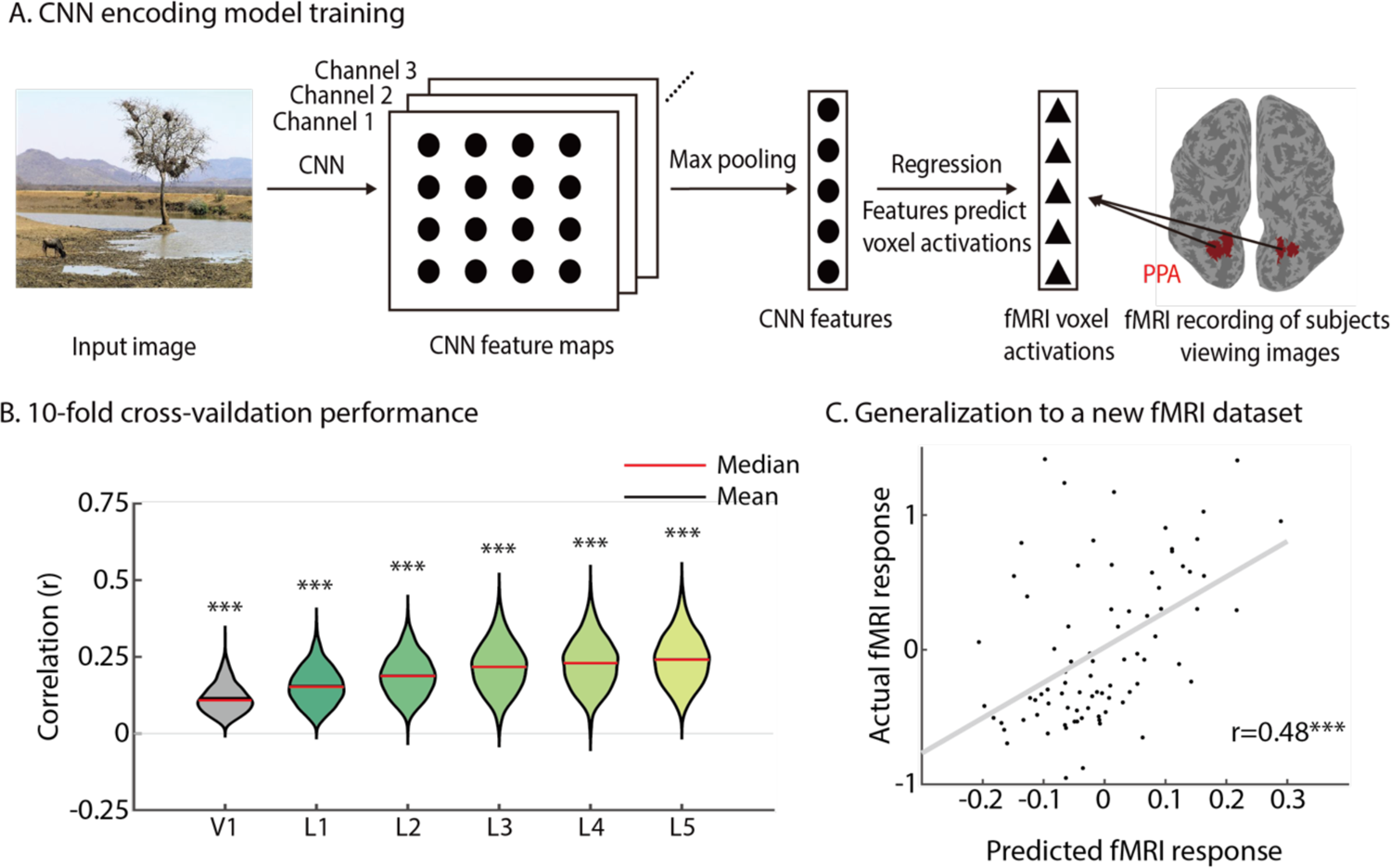
Encoding models and semantic preference mapping. **A)** Image-computable voxelwise encoding models of visual feature tuning were trained using the BOLD5000 fMRI dataset and the convolutional layers of a pre-trained AlexNet CNN. The encoding models were created by truncating AlexNet at a convolutional layer, adding a global max pooling operation, and then training a linear regression layer to map CNN feature activations to image-evoked fMRI responses. **B)** Encoding models were evaluated using a 10-fold cross-validation procedure. Performance generally increased along the layers of the CNN. For comparison, encoding models were also constructed using a computational model of V1 instead of a CNN. These violin plots show the distribution of encoding model performance across all voxels in all subjects for each ROI. The p-values were calculated by permutation tests (N=10,000 iterations). **C)** A strong test of generalization performance was conducted using data from Bonner & Epstein, 2021. The trained encoding models from BOLD5000 were used to generate predicted univariate fMRI responses in the PPA for a new set of stimuli and subjects. This plot shows the correlation between the predicted and actual fMRI responses in the PPA, which was strongly significant (r=0.48, p=1e-5). CNN: Convolutional neural network. PPA: parahippocampal place area, ***p<0.001.

We assessed the reliability of encoding model performance in two ways. We first used cross-validation to examine out-of-sample prediction accuracy on the BOLD5000 dataset and found that the prediction accuracy in the PPA was statistically significant (Fig. 1B, see Supplementary Fig. 2 for other ROIs). For our follow-up analyzes, we primarily focused on encoding models built from layer 5 because this layer showed the highest encoding model performance, but our results are not contingent on the specific convolutional layer used to build the encoding models, and our key findings are largely consistent across all convolutional layers (see, for example, Supplementary Figure 9A). We next performed a stringent test of how well our encoding models could generalize to new data by examining their ability to predict the average responses of the PPA in a separate fMRI dataset with new subjects and stimuli. We examined data from an fMRI study of object representation in which four subjects viewed images of isolated objects from 81 different categories^32^. We found that our encoding models trained on the BOLD5000 dataset were highly accurate at predicting the fMRI responses of the PPA to novel stimuli in a completely different set of subjects (Fig. 1C; r=0.48, p=1e-5). The strong generalization performance of our encoding models suggests that they capture important aspects of the feature preferences of the PPA and, furthermore, that these feature preferences explain a substantial portion of variance in the responses of the PPA to a wide range of stimuli, including complex scenes and isolated objects.

### Selectivity for scenes and object properties

With our computational model of PPA feature tuning in hand, we next sought to determine whether visual feature preferences in a feedforward model are sufficient to reproduce the multifaceted selectivity profile of the PPA for scenes, high-level object properties, and low-level contour statistics. We developed an approach that builds on a powerful two-fold procedure for characterizing cortical tuning profiles: first, highly parameterized encoding models are fit to neural data (as we have done with our computational encoding models), and second, *in silico* experiments are performed to reveal the interpretable, latent properties of these models^24, 33–35^. The strength of this two-fold procedure is that it combines the predictive power of highly parameterized models with the interpretability gained from *in silico* experiments. Specifically, we developed an approach for using high-throughput experiments to rigorously assess the latent information content of visual features in the context of a large natural scene dataset. In doing so, we are able to address a critical challenge for studies of visual feature coding: namely, that hierarchically computed visual features are notoriously difficult to understand through direct visualization methods but may nonetheless correspond to interpretable directions along the natural image manifold^36–38^. In fact, we attempted to directly visualize the image features that drive the responses of our PPA encoding models, but we found that these feature visualizations were largely inscrutable (Supplementary Fig. 3).

We first sought to determine whether our mid-level model of the PPA exhibits a pattern of scene-selectivity in its mean activation to images of scenes, objects, and faces, which are the stimulus categories that are often used to localize the PPA (Fig. 2A). In the following analyses, we refer to our computational encoding model of the PPA as simPPA (and we use a similar naming convention for other ROIs in the supplementary figures). We found that much like the actual PPA, the activations of our simPPA model to a set of localizer stimuli showed the typical pattern associated with scene-preferring areas, with response preferences ordered from scenes to objects to faces. In direct comparisons, the mean activation of simPPA was significantly greater for scenes relative to both objects (t(702)=22.00, p=1e-96) and faces (t(702)=56.95, p=1e-100) (Fig. 2B). It is worth noting that the scene-selectivity of simPPA is driven by visual feature tuning and cannot be attributed to spatial preferences, given that our encoding models involved a global max pooling procedure. Furthermore, although simPPA was trained to predict PPA responses to a diverse sample of natural scenes, it was never trained on the localizer stimuli examined here. Thus, these findings show that the visual feature tuning of simPPA is sufficient to generate a reliable pattern of scene-selectivity that generalizes to new stimuli.

**Figure 2.**
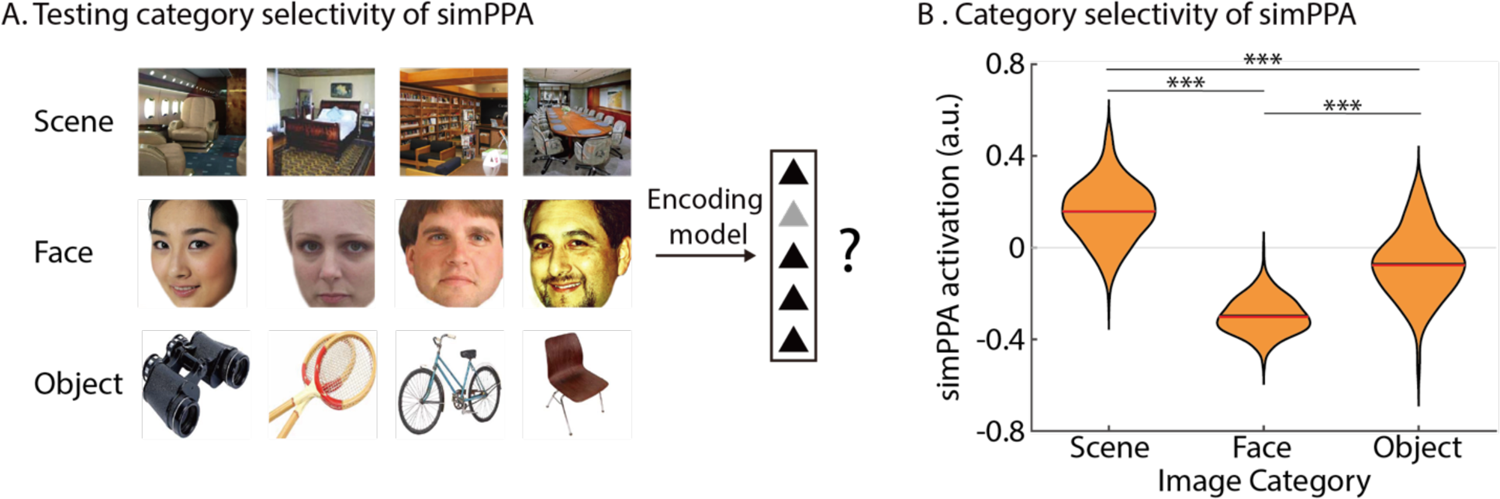
Visual feature tuning in simPPA gives rise to scene-selective responses. **A)** simPPA responses were obtained for a set of standard functional localizer images, including scenes, faces, and objects. **B)** simPPA showed preferential responses to scenes relative to both faces and objects. Red and black lines indicate the median and mean of the distributions. a.u.: arbitrary unit. ***p<0.001.

We next sought to characterize the selectivity of simPPA for object properties. We were specifically interested in determining whether the visual feature preferences of simPPA are reliably associated with interpretable object properties in the statistics of natural scenes. To accomplish this, we developed a computational method to characterize how the activations of an image-computable encoding model are affected by the presence of specific object categories in the context of natural images, and we used behavioral studies to relate these findings to human-interpretable object properties. This approach, which we refer to as semantic-preference mapping, has several strengths. First, it allows us to determine how the seemingly inscrutable feature representations of simPPA are related to the nameable components of scenes (i.e., objects). Second, it allows us to determine how simPPA responds to objects in their natural image contexts. And third, it is scalable to a large sample of images (i.e., 10^4), allowing us to characterize the association between visual features and interpretable object properties in a manner that is broadly representative of natural scene statistics.

Semantic-preference mapping works by systematically occluding instances of objects from target categories in a large set of images and then assessing how model activations are affected by the occlusion of these objects (Fig. 3A). We used the ADE20K dataset of densely annotated scenes to perform targeted occlusions of objects from specific semantic categories. The ADE20K dataset contains 27,574 images of real-world scenes from a diverse array of scene categories^39^. The objects in each image of this dataset have been manually segmented and labeled by an expert human annotator. We examined 85 categories of objects that each had at least 500 instances in the ADE20K dataset (these categories are listed in Supplementary Table 1). We performed targeted occlusions of all instances of these object categories and passed the occluded images to our encoding model (see Methods for details). For all units in the encoding model, we calculated the difference in activation for each occluded image relative to its corresponding original image, and we then calculated the mean of this difference score across all instances of an object category. The resulting metric indicates how strongly the responses of the encoding model are affected by the presence of a target object category in an image (Fig. 3A). We refer to this metric as a selectivity index. As an illustrative example, if a unit in the encoding model hypothetically responded to the features of cars, then its responses would decrease whenever cars were occluded, and it would have a high selectivity index for the target category *car*. Note that we partialled out occluder size from the selectivity indices to ensure that our results could not simply be attributed to differences in occluder size across categories (see Methods for details). We also performed several experiments that verified the robustness of the semantic-preference mapping results to variations in occluder shape (i.e., oval vs. rectangle) and CNN initialization parameters (see Methods and Supplementary Fig. 4).

**Figure 3.**
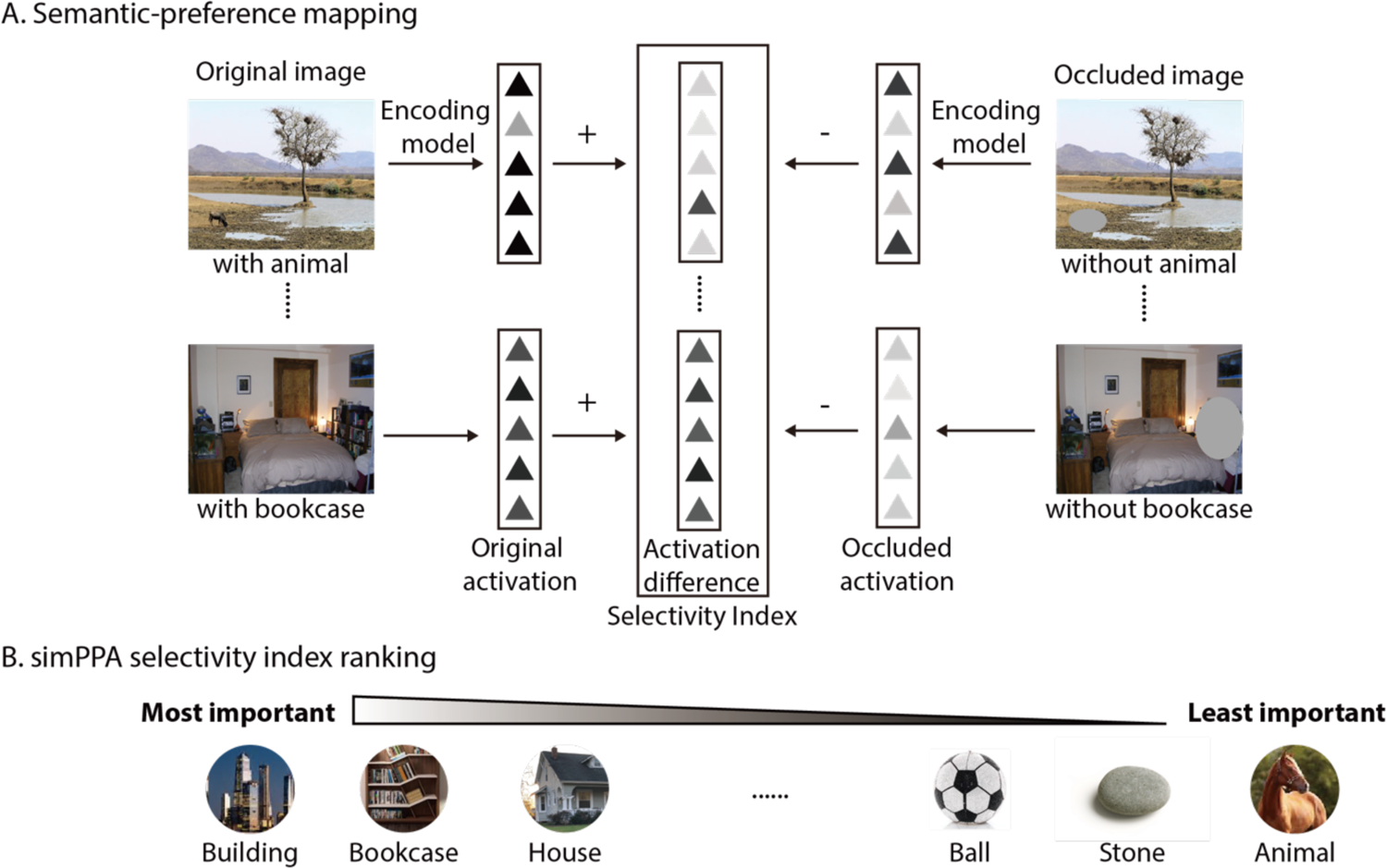
Semantic preference mapping. **A)** In the semantic preference mapping procedure, a database containing densely segmented images is used to perform targeted occlusions of object categories and to assess how encoding model activations are affected by these object occlusions. We used the ADE20K database for these analyses^39^. This procedure is repeated for all instances of an object category in the database, and the results are averaged to produce a selectivity index for each object category. **B)** This panel illustrates the results of the semantic preference mapping procedure for simPPA by showing the object categories with the highest and lowest selectivity indices.

After calculating the selectivity indices for all 85 object categories, we then performed follow-up experiments to determine whether these selectivity indices were related to human-interpretable object properties. Specifically, we collected behavioral ratings for five properties that have previously been linked to the responses of the PPA: large in real-world size, fixed in location, inanimate, manmade, and rectilinear^13, 16, 19, 40^ (Fig. 4A and Supplementary Figs. 5 and 6; see Methods for details). Because the manmade and inanimate ratings were highly correlated (r=0.91), we combined them into a single rating by taking their averaging for each category. We then calculated correlations of these object property ratings with the selectivity indices from our semantic-preference mapping procedure. For these correlations, we partialled out the size of the occluder for each object category to ensure that the correlations could not be attributed to occluder size (see Methods). We first calculated correlations with the mean selectivity index across all units in simPPA, which is analogous to examining the global univariate response of a brain region.

**Figure 4.**
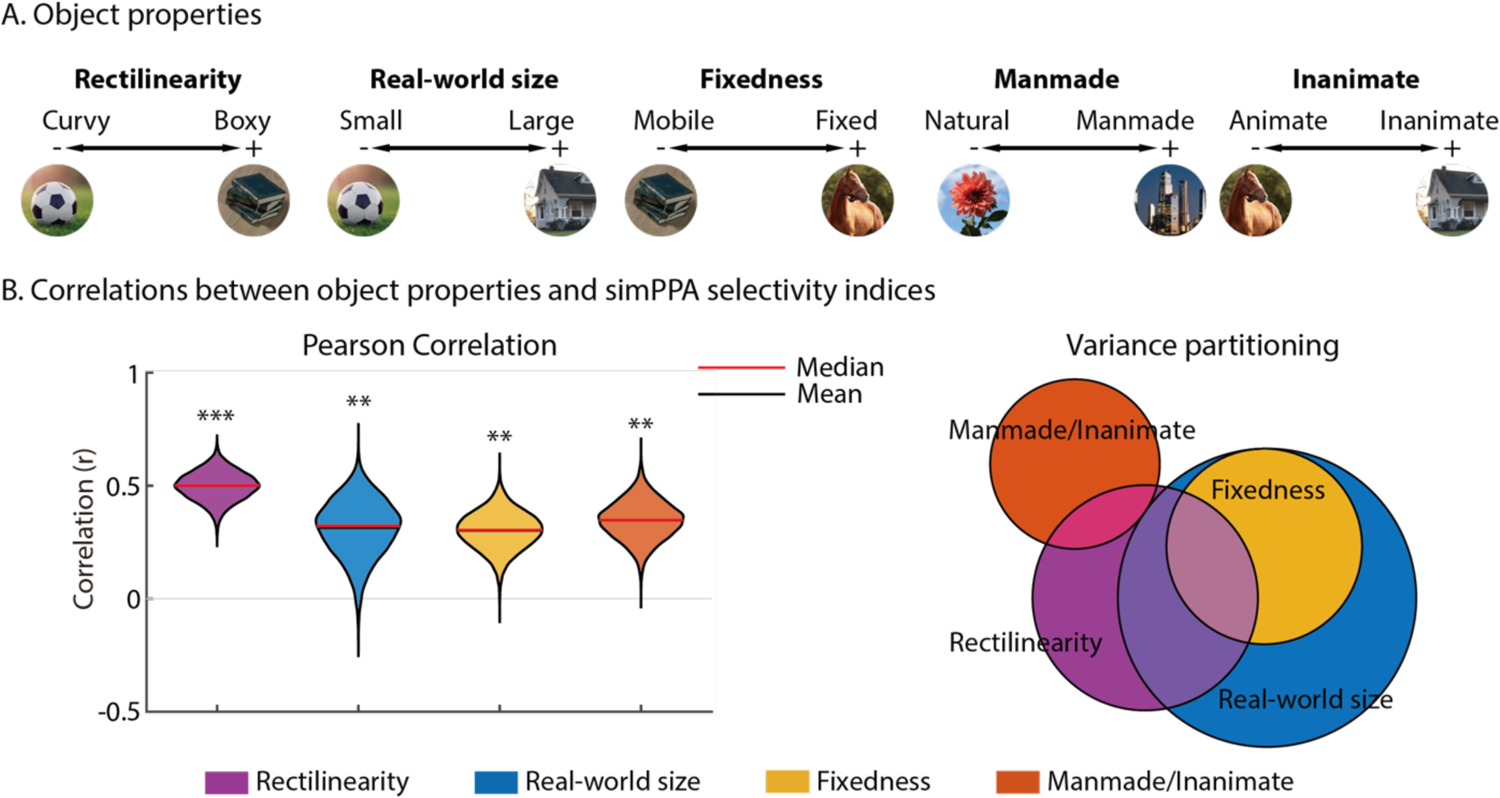
Visual feature tuning in simPPA gives rise to preferential responses to multiple high-level object properties. **A)** Behavioral ratings were collected for object properties that have previously been associated with the responses of the PPA: rectilinearity, real-world size, fixedness, manmade, and inanimate. These ratings were collected for all 85 object categories that were examined in the semantic preference mapping procedure. Because the manmade and inanimate ratings were highly correlated, they were averaged to create a single manmade/inanimate rating. **B)** The average selectivity indices of simPPA were significantly correlated with all four object properties (left). This shows that the visual feature tuning of simPPA gives rise to preferential responses to objects that are rectilinear, large in real-world size, fixed in location, and inanimate/manmade. The violin plots show distributions of the correlation values across 10,000 bootstrap resampling iterations. Venn diagram (right) generated by variance partitioning was used to identify the unique and shared contributions of each object property for explaining variance in the selectivity indices of simPPA. There was a considerable amount of shared variance across the object properties. However, all properties other than fixedness also explained some unique variance. **p<0.01, ***p<0.001. All p-values were calculated by permutation tests (N=10,000 iterations).

**Figure 5.**
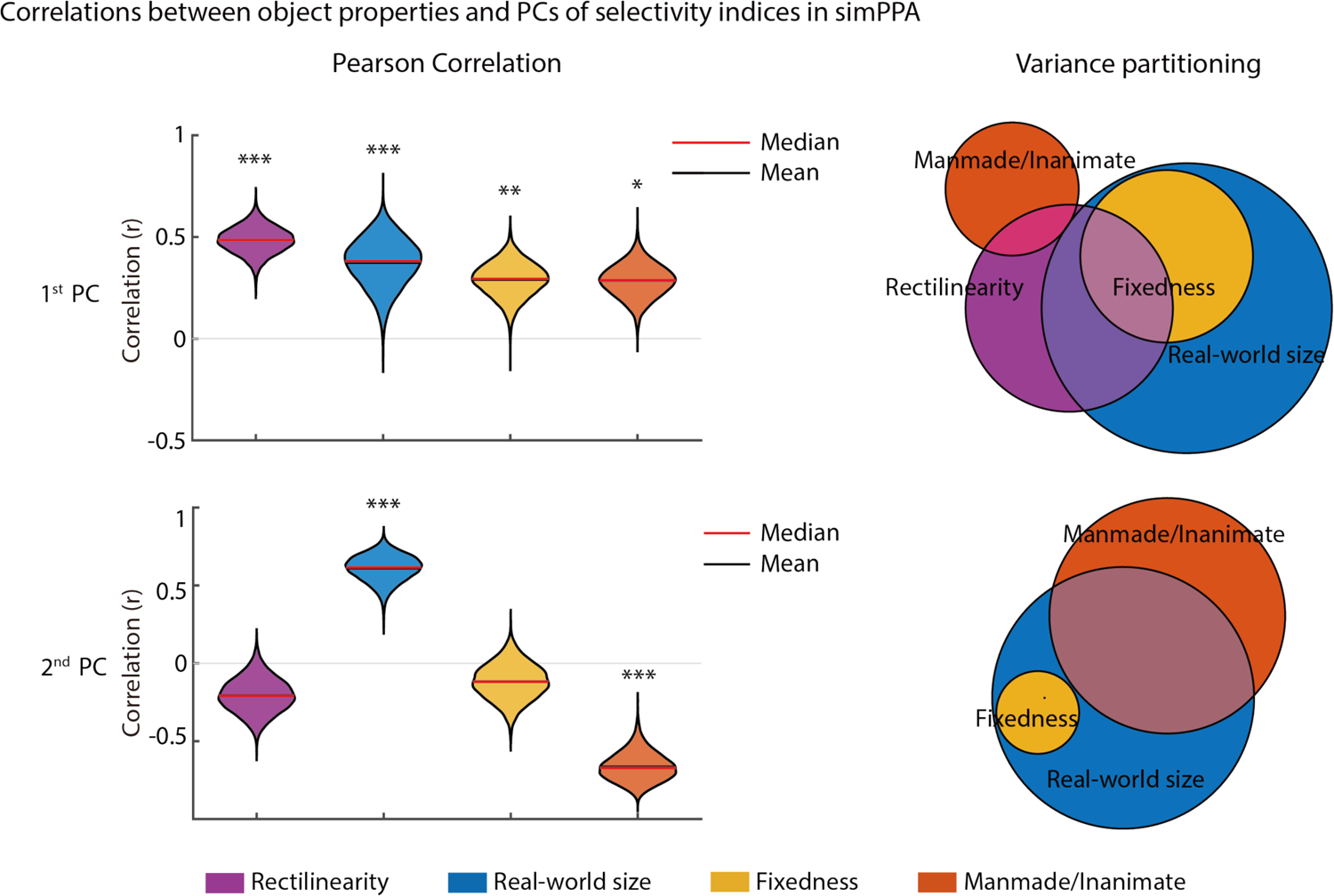
Visual feature tuning in simPPA gives rise to orthogonal dimensions of high-level object preferences. As shown on the top left, the first PC of the selectivity indices from simPPA was significantly correlated with all four object properties and resembled the findings for the global univariate selectivity indices in Figure 4. As shown on the bottom left, the second PC of the selectivity indices exhibited a different pattern. This PC had significant but opposite-signed correlations with real-world size and manmade/inanimate and appears to reflect a preference for large, natural objects. The violin plots show distributions of the correlation values across 10,000 bootstrap resampling iterations. Variance partitioning of both PCs were used to identify the unique and shared contributions of each object property for explaining variance in the selectivity indices of simPPA. For both PCs, there was a considerable amount of shared variance across the object properties. For the first PC, there are unique contributions from all properties other than fixedness. For the second PC, only inanimate/manmade and real-world size had unique contributions. *p<0.05, **p<0.01, ***p<0.001. All p-values were calculated by permutation tests (N=10,000 iterations). PC: Principal component.

**Figure 6.**
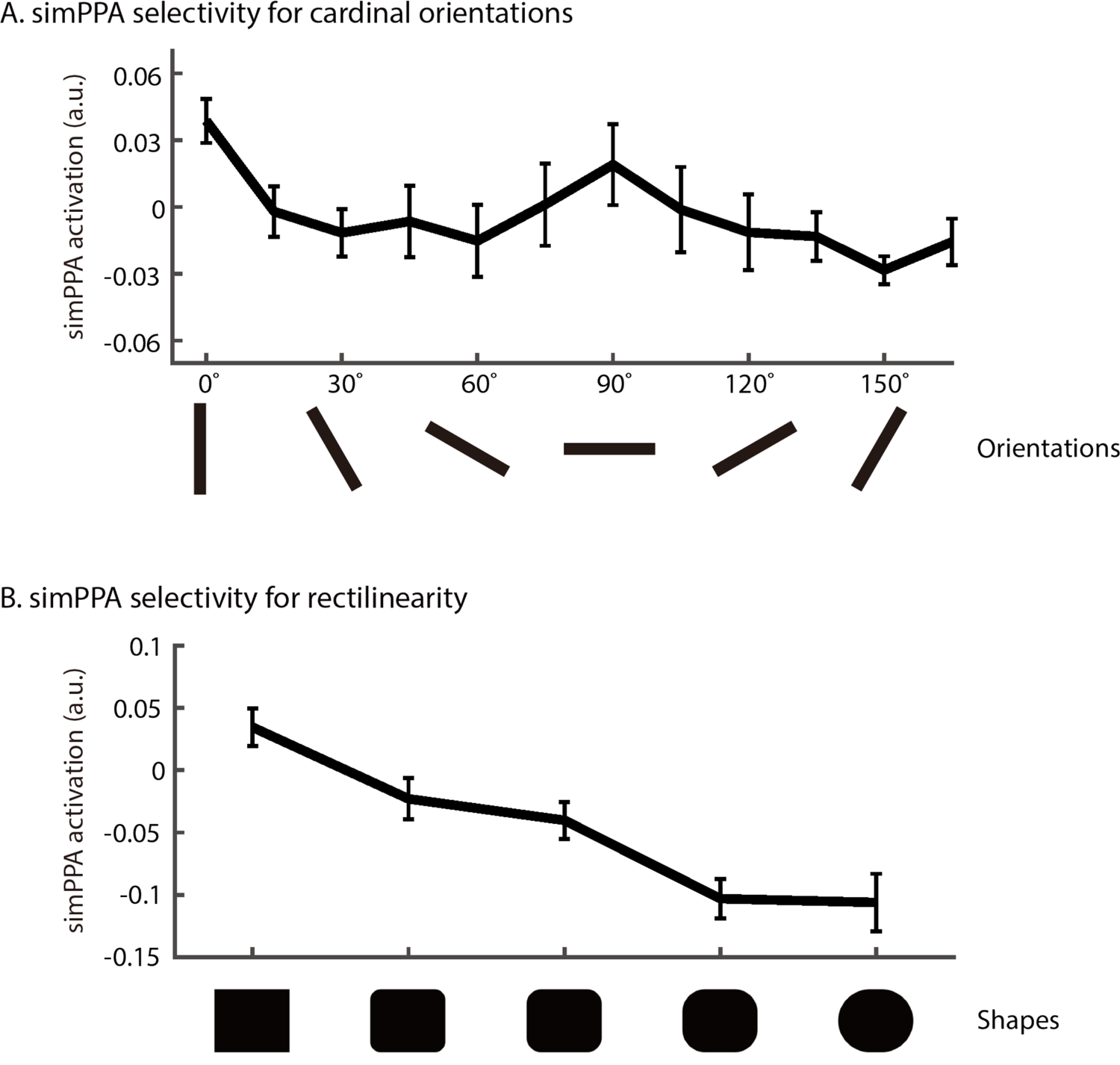
Visual feature tuning in simPPA gives rise to preferential responses to cardinal orientations and rectilinear shapes. The selectivity of simPPA for low-level perceptual properties was assessed using minimal stimuli containing oriented Gabor patches or simple shapes. A) The average univariate response of simPPA is plotted for stimuli containing Gabor patches at a range of angles from 0° to 165°. Error bars represent +/-1 SD across the units of simPPA. These findings show that simPPA responds more to contours at cardinal orientations (0° and 90°). B) The average univariate response of simPPA is plotted for stimuli containing simple shapes that varied along a continuum from boxy to curvy. Error bars represent +/-1 SD across the units of simPPA. These findings show that simPPA responds more to rectilinear shapes.

We found that the mean selectivity index was significantly correlated with all four object properties, indicating that simPPA exhibits a preference for objects that are boxy, large in real-world size, fixed in location, and inanimate/manmade (Fig. 4B and Supplementary Figs. 7 & 8A). By examining other CNNs (including CNNs pre-trained on different image sets) and CNN layers, we found that these correlations were not a fluke of the specific CNN encoding models used for our primary analyses but, rather, reflected robust correlations between high-level object properties and surprisingly simple feedforward visual features in CNNs.

Indeed, we observed similarly strong correlations for all four object properties when examining PPA encoding models built from other CNN layers, other CNN architectures, and another CNN pre-training dataset (Supplementary Figure 9 and Supplementary Table 2). In contrast, when examining PPA encoding models built from a V1-like computational backbone, we found that real-world size was the only property with a similarly strong correlation to our CNN-based models (Supplementary Figure 8). All other object properties had significantly stronger correlations with the CNN-based models, suggesting that the relevant visual features for these effects reflect a level of complexity that goes beyond the basic edge detection of a V1-like model. These results demonstrate that the known object preferences of the PPA, even for seemingly high-level properties like real-world size and fixedness, can emerge from purely feedforward computations of visual features and that these effects are representative of the statistical regularities in a large and diverse sample of natural scenes.

We next performed variance partitioning analyses to determine the degree to which our object-property ratings accounted for unique and shared variance in the selectivity of simPPA (see Methods for details). Rectilinearity had the highest correlation with the mean selectivity of simPPA (Fig. 4B left), and our variance partitioning analyses showed that it accounted for a portion of the explained variance associated with all three other object properties (Fig. 4B right). For fixedness, the explained variance could be fully accounted for by real-world size. However, real-world size, manmade/inanimate, and rectilinearity had unique explained variance that could not be attributed to any other property (Fig. 4B). Thus, the mean response of simPPA exhibits preferences for the object properties of rectilinearity, real-world size, and manmade/inanimate that cannot be fully reduced to a common underlying factor.

Our analyses thus far have focused on the overall mean selectivity of simPPA. However, it is possible that these selectivity indices contain multiple latent dimensions of object preferences. We next performed analyses to examine the multivariate selectivity profile of simPPA and its principal representational dimensions. We applied principal component analysis (PCA) to the selectivity indices of simPPA for all 85 object categories from the semantic-preference mapping procedure. We focused on the first two principal components (PCs), which accounted for 83.4% and 11.3% of the variance in the selectivity indices. We then analyzed these PCs in the same way as the mean selectivity index. The first PC largely resembled the mean selectivity index, with significant correlations with all four object properties and unique explained variance for every property except fixedness (Fig. 5 and Supplementary Fig. 8B). Though the second PC accounted for far less variance than the first PC, it exhibited an interesting pattern of selectivity for large, natural/animate objects, with significant but opposite-signed correlations for real-world size and manmade/inanimate, which both explained unique variance (Fig. 5 right). Furthermore, there was almost no correlation with rectilinearity in the second PC. The results of these PC analyses show that when the multivariate selectivity of simPPA is broken down into its principal latent dimensions, we find two orthogonal patterns of selectivity: one for objects that are large and manmade and another for objects that are large and natural.

### Selectivity for cardinal orientations and rectilinear shapes

One of the most perplexing aspects of the PPA is that in addition to its selectivity for scenes and high-level object properties, it also exhibits preferential responses to low-level geometric stimuli with a high proportion of rectilinear shapes and cardinal orientations^18–20^. Here we tested whether our simPPA model of visual feature tuning also exhibits a similar pattern of response preferences for simple geometric stimuli. We created two sets of simple low-level stimuli to examine the response profile of simPPA across contour orientations and degrees of rectilinearity. We first examined simPPA responses to minimal images containing a single Gabor patch at a specific orientation, ranging from 0 to 165 degrees in 15-degree intervals (Fig. 6A). We found that, as in previous reports of the PPA, simPPA shows a response preference for contours at cardinal orientations (i.e., vertical and horizontal). An analysis of other ROIs showed that this preference for cardinal orientations was not a universal phenomenon of our encoding models for all ROIs (Supplementary Fig. 10). We next examined simPPA responses to minimal images containing simple shapes that varied along a continuum from curvilinear to rectilinear (Fig. 6B). Again, much like previous reports of the PPA, simPPA showed a response preference to simple geometric stimuli with rectilinear features. This preference for rectilinear features was not a universal phenomenon of our encoding models for all ROIs (Supplementary Fig. 11). Together, these findings show that the feedforward computation of visual features in simPPA gives rise to a multifaceted selectivity profile for scenes and object properties in natural images as well as preferences for low-level contour statistics in minimal stimuli.

## DISCUSSION

We fit a feedforward model of visual feature tuning to the scene-evoked fMRI responses of the PPA and used high-throughput experiments to show that it reproduces core aspects of PPA selectivity for scenes, object properties, and simple geometric stimuli. Our results provide a unified theoretical account that resolves several seemingly divergent findings on the response preferences of the PPA, and they show how tuning for surprisingly simple visual features can give rise to rich patterns of selectivity that span multiple levels of stimulus complexity.

Our results have implications for understanding the organizing principles of the ventral visual stream. One of the central anatomic properties of the ventral stream is its organization into patches that are selective for categories, such as places, faces, and objects, and coarse conceptual domains, such as those based on animacy and real-world size^40–42^.

Although this functional organization of the ventral stream has long been characterized in terms of high-level, interpretable stimulus attributes, such as categories, recent findings suggest that the fundamental organizing principles may be better characterized in terms of differential tuning to visual features^25, 36, 37^. One such finding showed that mid-level visual features are sufficient to elicit domain-selective fMRI activations in the ventral stream for the properties of animacy and real-world size, even when the experimental stimuli are unrecognizable as natural objects^37^. Another key finding showed that the category-selective organization of object representations in the macaque ventral stream can be mapped onto the first two principal components of a feedforward CNN and may thus naturally arise from the statistical structure of visual feature representations^36^. It has also recently been shown the feedforward neural network models provide more accurate predictions of response preferences in category-selective visual regions than descriptive models of image categories or the predictions of experts in the field^25^. More broadly, multiple studies have found that the representations of the human ventral stream are better explained by perceptual features than by the abstract properties that underlie category identity or human intuitions about semantic similarity^25, 29, 43–45^. Our findings are broadly consistent with a feature-based theory of ventral-stream organization and show the surprising degree to which feedforward models of visual feature tuning can account for the characteristic selectivity profile of the PPA for stimulus properties spanning from high-level, semantic attributes to low-level contour statistics.

Although hierarchically computed visual features are critical for explaining the representations of visual cortex, they are notoriously difficult to characterize^38^. We lack simple algorithmic models of mid-level features, in contrast to the Gabor model for V1. The most effective approach for discovering mid-level visual features that are predictive of cortical responses is deep learning in CNNs^28, 46^. However, the resulting CNNs are black boxes whose mid-level representations are challenging to visualize and even more challenging to describe in words—their features exist in an ineffable valley between the describable patterns of low-level vision (e.g., edges) and the intuitive concepts of visual semantics (e.g., objects). Here we sought to gain a more informative view of mid-level features by characterizing their covariance with nameable scene elements—a procedure we call semantic-preference mapping. This approach allowed us to combine the strengths of a CNN with the interpretability of a tuning profile across a set of object categories. Using this approach, we found that the visual features of our simPPA encoding model had latent covariance relationships with interpretable object properties and that these covariance findings were representative of the statistical regularities in the large and diverse sample of natural images examined here. These analyses provide a new perspective on PPA selectivity: they demonstrate that the strong responses of the PPA to scenes and landmark-like objects could in principle be mediated by the feedforward computation of visual features that covary with scenes and landmarks in the natural statistics of images.

It is important to point out that our findings do not simply reveal a confound between visual features and the high-level properties of scenes and landmarks. Rather, they reveal a potential mechanism for mediating the selectivity of the PPA during the passive viewing of natural images. After all, the PPA is a visual region that receives a large portion of its inputs from the visual pathway starting at the retina^47^. Any mechanistic theory of the PPA will ultimately need to explain how it processes these upstream inputs in a manner that yields rapid and automatic selectivity for scenes and object properties. As an analogy, we could consider V1 cells, which are commonly described as being functionally selective for edges and are mechanistically modeled using oriented and localized spatial-frequency filters^48^. The relationship between the mechanistic implementation (i.e., oriented spatial-frequency filters) and the functional selectivity (i.e., edges) is premised on the covariance between the filter responses and the presence of edges in images, but it does not require that this relationship be one of perfect mutual information. In fact, oriented spatial-frequency filters also provide information about image features other than edges and can even arise in models trained on spatially smooth images that contain no edges whatsoever^49^. Despite this, there is little disagreement that V1 can be functionally described in terms of edge representation and that the underlying computational mechanisms involve spatial-frequency filters. Similarly, we argue that the PPA can be functionally described as representing scene and landmark-like objects, and that one of the underlying mechanisms that directly supports this function is the feedforward computation of simple visual features.

It is also important to point out that our feedforward model does not capture all aspects of information processing in the PPA. Our model is only intended to account for the initial feedforward activations of the PPA and does not contain feedback and recurrent processes, which are pervasive in visual cortex and likely play a crucial role in the PPA. In fact, it is known that the PPA shows scene-related activation even without visual stimulation, including in subjects who are congenitally blind and in sighted subjects who are haptically exploring miniature scenes^22, 23^. Thus, there appear to be scene-specific top-down feedback mechanisms in the PPA that remain to be explained. Our model is also spatially coarse and is focused on capturing tuning for visual features rather than spatial receptive-field biases.

Although receptive-field biases are known to exist in the PPA^47^, we found that our spatially coarse encoding model could nonetheless account for a substantial portion of the global univariate response profile of the PPA. Future work could examine how tuning for visual features interacts with the receptive-field biases of the PPA and to determine whether there exist relevant covariance relationships between visual features and receptive-field locations in the natural statistics of vision^50^. Our model is also not intended to explain the effects of navigational experience on the PPA, which shows stronger responses to objects that occur at navigationally important locations and treats stimuli as more similar if they come from the same place^14, 15, 51^. Furthermore, our findings do not account for functional differences along the anterior-posterior extent of the PPA, which appear to reflect a general trend toward visual-form representations in the posterior PPA and mnemonic representations in the anterior PPA^32, 52, 53^. However, future studies could leverage our modeling framework to test whether the effects of navigational experience in the PPA involve the modulation of visual features from navigationally important stimuli and to test whether the mnemonic representations of anterior PPA are implemented through the associative coding of visual features from co-occurring stimuli, including object categories in scenes and distinct views of places^14, 32^.

Feedforward visual features may be the currency of the ventral stream^25, 36, 37^, and approaches for making sense of these features are crucial for advancing our understanding of visual cortex. Here we show that when computational models of feedforward feature tuning in visual cortex are combined with methods for revealing their interpretable properties, these methods reveal how cortical selectivity profiles naturally span multiple levels of stimulus complexity and they provide insight into the category-selective organization of the ventral stream. More broadly, our computational modeling framework paves the way for examining the behavioral significance of visual features in scene- and landmark-processing and for exerting control over representational states in the cortical scene network through targeted visual stimulation^26, 27^.

## MATERIALS AND METHODS

### fMRI data processing

We analysed data from the publicly available BOLD5000 dataset (https://bold5000.github.io), which contains 3T BOLD fMRI data in four subjects who viewed between 2,952 and 4,916 unique natural scene images depicting real-world environments and objects^30^. This dataset was designed to sample fMRI activations to a large and diverse set of natural scenes. To maximize stimulus diversity, most images in this dataset were presented a single time over the course of the experiment. Stimuli were presented for 1 sec followed by a 9 sec interstimulus interval. In the scanner, subjects performed a valence judgment task, responding with how much they liked the image using the metric: “like”, “neutral”, “dislike”. See^30^ for a more detailed description of this dataset.

All functional data were preprocessed using fMRIPrep^54^, which performed 3D motion correction, distortion correction, and co-registration to the T1 anatomical image. After preprocessing, we estimated the activation to each image using a series of general linear models that included a single regressor for each trial and another regressor for all other trials. This procedure has been shown to be more accurate for estimating activation magnitudes in event-related designs with high signal to noise^55^. We implemented this general linear modeling procedure using the function 3DLSS in AFNI^56^. These activation estimates were used as the predictands for our CNN encoding models.

ROIs were identified using four localizer runs. First, a group-based parcel derived from a large number of subjects was warped to each subject’s native space to act as an anatomical constraint^57^. Bilateral ROIs were identified within the parcel in each hemisphere by identifying the top 200 most activated voxels from the localizer contrast. PPA, OPA and RSC were identified using the scenes > objects contrast, LOC was identified using the objects > scenes contrast, and EVC was identified using the objects > scrambled contrast. In total, each subject had 400 voxels in each ROI.

### Encoding models

For our primary analyses, we constructed voxelwise encoding models using AlexNet pre-trained on Places^31^. Our modelling procedure involved pooling and reweighting of the CNN responses from a convolutional layer (after ReLU) to predict the image activation estimates from BOLD5000 (Fig. 1A). We applied global max pooling to obtain a single activation for each feature channel, and we passed these feature activations to a linear regression layer that was trained to predict the image-evoked fMRI activations as a weighted sum of the CNN feature activations. We trained the linear regression layer using LASSO (L1 penalized) regularization. A 10-fold cross validation procedure was used to search for the optimal regularization penalty in each voxel. The penalty parameter was selected from 20 values on a log-scale from 1e-3 to 1e5. After identifying the optimal penalty parameter for each voxel, we learned a set of regression weights using this penalty parameter and the full set of fMRI data. Together, the truncated CNN, followed by max pooling, and the regression layer define an image-computable encoding model of mid-level feature tuning for each voxel.

We were interested in L1 regularization as a means of learning sparse encoding models that emphasize the CNN features that are most important for each voxel. However, we were unsure if L1-regularized regression would perform as well as L2-regularized or ordinary least squares (OLS) regression. We therefore evaluated the performance of different regression methods by running encoding-model analyses on the BOLD5000 dataset with 10-fold cross-validation using OLS regression (without regularization), LASSO regression (L1 regularization) and ridge regression (L2 regularization). Although previous studies have typically used ridge regression when fitting voxelwise encoding models^33, 34^, we found that LASSO outperformed both ridge and OLS (Supplementary Fig. 1). Thus, our encoding models benefited from a sparse regularization procedure that pushes some of the regression weights to zero and emphasizes the subset of feature activations that are most informative for each voxel.

Encoding model performance was evaluated in two ways. First, we performed a new 10-fold cross-validation procedure on the BOLD5000 dataset while keeping the regularization penalty fixed (using the previously learned optimal penalty for each voxel). The cross-validation scheme used for this evaluation was different from the cross-validation scheme that was used when selecting the regularization penalty. Supplementary Fig. 2 shows the mean Pearson correlations between the predicted activations and the observed activations across all cross-validation folds and all voxels in each ROI. Note that because the penalty parameter was learned on the same data, the performance estimates may be biased upwards. We therefore performed an additional stringent evaluation of encoding-model generalization performance using a separate set of fMRI data with new subjects and new stimuli, which is described below. It is also worth noting that the encoding models perform well above chance in the BOLD5000 dataset even when using OLS regression without regularization, which means that regularization is not required to achieve statistically significant performance. Furthermore, our results and conclusions do not depend on the specific values of the performance estimates in BOLD5000. It is already well-established that CNNs are state-of-the-art encoding models of fMRI responses in visual cortex^28^. The primary goal of our analyses is to characterize the mid-level representations of these encoding models after they have been fit to fMRI data.

Second, to rigorously test generalization performance, we used the trained encoding models from the BOLD5000 dataset to predict the fMRI activations to 81 object categories from a separate fMRI dataset with a different set of subjects. We used the fMRI data from Bonner & Epstein, 2021 (https://osf.io/ug5zd/), which included fMRI responses in four subjects who viewed images of isolated real-world objects from 81 different categories that were presented on meaningless textured backgrounds. In the scanner, the subjects performed a simple oddball-detection task of pressing a button whenever a warped object was shown.

See the original publication for a detailed description of these data^32^. Each object category in this dataset contained 10 unique images, which were shown in a block design. We ran all images through our CNN encoding models and obtained the average activation across all 10 images for each object category. Our goal was to test whether these encoding model activations could predict the average univariate fMRI activation of our ROIs. For each ROI, we averaged the encoding model activations across all voxels in all subjects from BOLD5000 to obtain a single activation value for each object category, which we compared with the actual univariate fMRI activations averaged over all subjects in the Bonner & Epstein data. We observed a strong correlation between the predicted activations from our encoding models and the actual fMRI activations in PPA (Fig. 1C). These findings demonstrate that the encoding models trained on the BOLD5000 dataset exhibit remarkable generalization performance across both subjects and stimuli when predicting the univariate activations of multiple ROIs (including PPA). Thus, our encoding models appear to capture key aspects of the mid-level feature tuning in these ROIs.

### CNN feature visualization

To visualize the representations encoded in a CNN units, we utilized a CNN feature visualization technique^58^. Since the goal of this analysis was to visualize units that were important in predicting PPA responses, we first sorted the units based on their average weighting in simPPA encoding model. Then for the first nine units with the highest average weighting, we generated feature visualizations. To perform the visualization process, an input image was randomly initialized, and we used the Adam optimizer to learn pixel values that strongly activate the targeted CNN unit. We ran 70 iterations of the optimization process. See Supplementary Fig. 3 for results.

### Category selectivity

To test whether simPPA exhibits classic category selectivity pattern, classic fMRI localizer stimuli, including scene, face and object images were passed through simPPA to obtain its activation (Fig. 2A). For each category, 352 images were used. Activations of each image were then averaged across the simulated voxels, then the activation distribution of images across categories were plotted in Fig. 2B.

### Semantic preference mapping

We developed an algorithmic approach to examine how the activations of our CNN encoding models were affected by the object classes present in an image. For this procedure, we made use of the ADE20K dataset, which contains 27,574 images of real-world scenes from a diverse array of scene categories^39^. The objects in each image of this dataset have been manually segmented and labeled by an expert human annotator. We used these segmentation masks to perform targeted occlusions of objects in images and assess how these occlusions affected the activation of the CNN encoding models (Fig. 3A). The logic of this procedure is that if an encoding model preferentially responds to certain categories of objects, then its responses will be strongly affected by occlusions of those objects. Our goal was to rigorously assess how the CNN encoding model activations were affected by the presence of these object categories in a large sample of images. We therefore examined all object categories that had at least 500 instances in the ADE20K dataset, which yielded a total of 85 categories (these are listed in Supplementary Table 1). For our targeted occlusions, we used the object segmentations to create the smallest oval mask that covered the target object. These masks contained random RGB values in each pixel, and the edges of these masks were blurred by morphological dilation using the Matlab function imdilate. We passed the occluded images to our CNN encoding models and calculated a difference score by subtracting the activation to the occluded image from the activation to the corresponding original images (without occlusion). We then calculated the mean of this difference score across all instances of an object category. The resulting metric indicates how strongly the responses of the encoding model are affected by the presence of a target object category in an image. We refer to this metric as a selectivity index. To ensure that our findings could not simply be attributed to the size of the occluders, we partialled out occluder size by regressing the selectivity indices against occluder size (i.e., mean number of pixels) and retaining the residuals, which we used for all follow-up analyses. For univariate analyses of each simROI, we averaged the selectivity indices across all voxelwise models in all subjects. When performing PCA for each simROI, we concatenated the selectivity indices across all voxelwise models in all subjects.

We performed analyses to assess the robustness of the results obtained from the semantic preference mapping procedure. We first ensured that our findings were not contingent on the specific shape of the occluder (i.e., oval) by repeating our analyses using rectangular occluders. We found that the mean selectivity indices in each simROI were highly consistent whether we used oval occluders or rectangular occluders (Supplementary Fig. 4; all r-values >0.65, all p-values<1e-5). We next evaluated whether the results of the semantic preference mapping procedure were consistent when using CNNs with different random initializations during pretraining. To do this, we examined 10 different instances of AlexNet trained on the CIFAR dataset using different random initializations^59^ (https://osf.io/3xupm/). We performed our entire pipeline of training encoding models and performing semantic preference mapping using these 10 different instances of AlexNet, and we compared the resulting selectivity indices across all 10 instances. We found that the mean selectivity indices in each simROI were highly consistent across all 10 instances of AlexNet (the mean pairwise correlations were greater than 0.9 for all simROIs).

### Behavioral ratings of object properties

Fifty subjects were recruited online through the Prolific platform. This experiment was in compliance with procedures approved by the Johns Hopkins University Institutional Review Board. Subjects were asked to judge five object properties for a highlighted object in an image using a 7-point scale (Supplementary Fig. 5). The judged object properties included curvature, real-world size, inanimate, manmade and fixedness. Each subject was presented with one image per each of the 85 object categories, with a total of 85 stimuli per subject. Stimuli were randomly chosen from the images used in the semantic preference mapping procedure. Subjects had the option of hovering a virtual magnifying glass over the image to enlarge any part of the image that was not clear. For each property, we used the average rating across all subjects in all follow-up analyses. As expected, some of these ratings covaried (Supplementary Fig. 6). We found that inanimate and manmade were highly correlated (r=0.91, p=1e-6), and we therefore decided to take the average of these two properties to create a combined manmade/inanimate rating.

### Variance partitioning

We used variance partitioning to evaluate the degree to which the object-property ratings explained unique or overlapping variance in the selectivity indices. We performed these analyses using the vegan package in R^60^. In these analyses, multiple object properties were used to predict the selectivity indices. Through a series of regressions using different subsets of object properties, we obtained the unique and shared variance associated with all object properties (see Figs. 4 and 5).

### Analyses of contour orientations and rectilinear shapes

To test whether the encoding models preferred contours at specific orientations, we created minimal images with Gabor patches at different orientations, ranging from 0° to 165° in 15° intervals (see Fig. 6A and Supplementary Fig. 10). These images were 492-by-402 pixels in size and contained a single Gabor patch in the center that has a wavelength of 100 (100 pixels/cycle) with spatial frequency bandwidth of 1 and the spatial aspect ratio of 0.5. We also evaluated encoding model responses to minimal images containing simple geometric shapes. We created a series of stimuli that varied along a continuum from boxy to curvy (see Fig. 6B and Supplementary Fig. 11). These images were 720-by-720 pixels in size and contained a single shape in the center that spanned ∼385 pixels in height and ∼460 in width.

### Analyses of encoding models with other computational backbones

We examined voxelwise encoding models of the PPA using different computational backbones, including AlexNet trained on Places^31^, AlexNet trained on ImageNet^61^, VggNet 16 trained on ImageNet^62^, Resnet 18 trained on Places^31^, Resnet 18 trained on ImageNet^63^ and DenseNet trained on Places^31^. In addition, a V1 computational model was also used to construct voxelwise encoding models of the PPA^64^. The V1 model contained Gabor filters in four different orientations and six different sizes/wavelengths whose outputs were passed to a half-wave squaring nonlinearity^64^. The output vector of the V1 model was treated in the same way as the CNN feature vectors. We constructed encoding models using the same procedures as those used for the Places-trained AlexNet model in our primary analyses, and we used semantic preference mapping to examine how the selectivity indices of these various computational encoding models correlated with the object property ratings (see results in Supplementary Figs. 9B & 9C and Supplementary Table 2).

## ACKNOWLEDGMENTS

We thank Alon Hafri and Emalie McMahon for comments on this manuscript. The data from Bonner & Epstein 2021 was funded by NIH grant R01EY022350.

## SUPPLEMENTARY INFORMATION

### Supplementary figures and tables

**Supplementary Figure 1.**
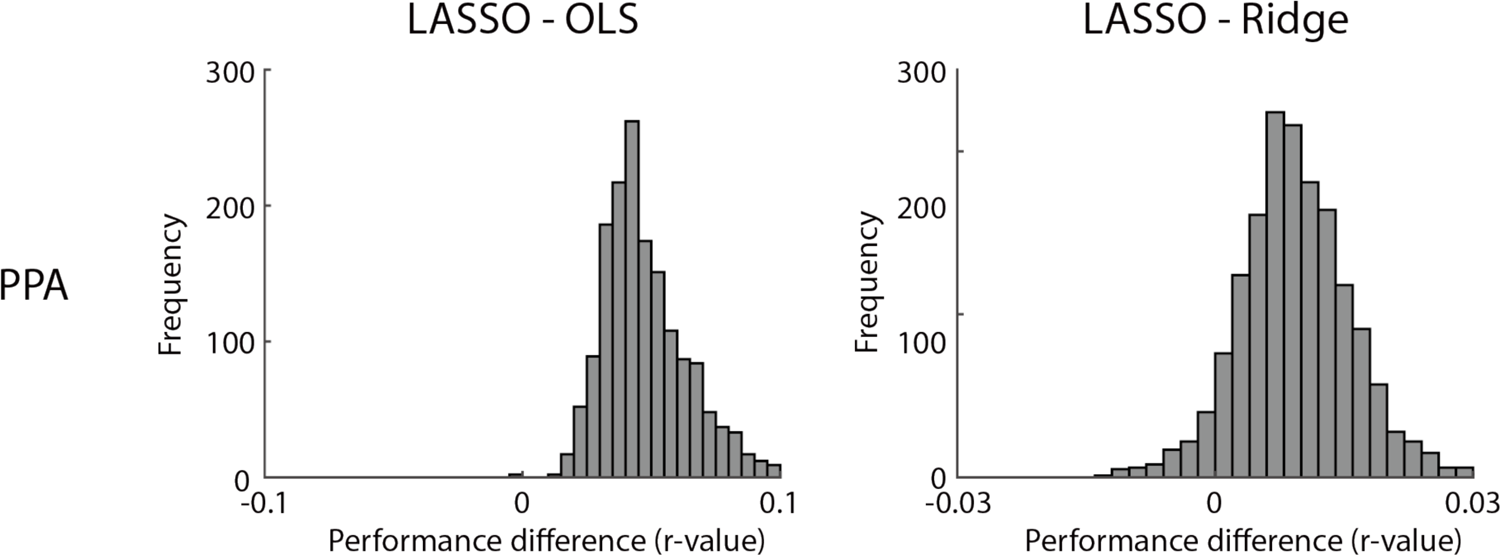
Distribution of performance comparisons between regression methods. These plots show distributions of difference scores between the cross-validated prediction accuracies of voxelwise encoding models in the PPA that were trained using different regression methods. The difference scores were calculated by subtracting the prediction accuracy when using OLS (left panel) or ridge (right panel) from the prediction accuracy when using LASSO. These plots show that LASSO outperformed OLS and ridge in nearly all voxels. OLS: Ordinary least square. LASSO: Least absolute shrinkage and selection operator.

**Supplementary Figure 2.**
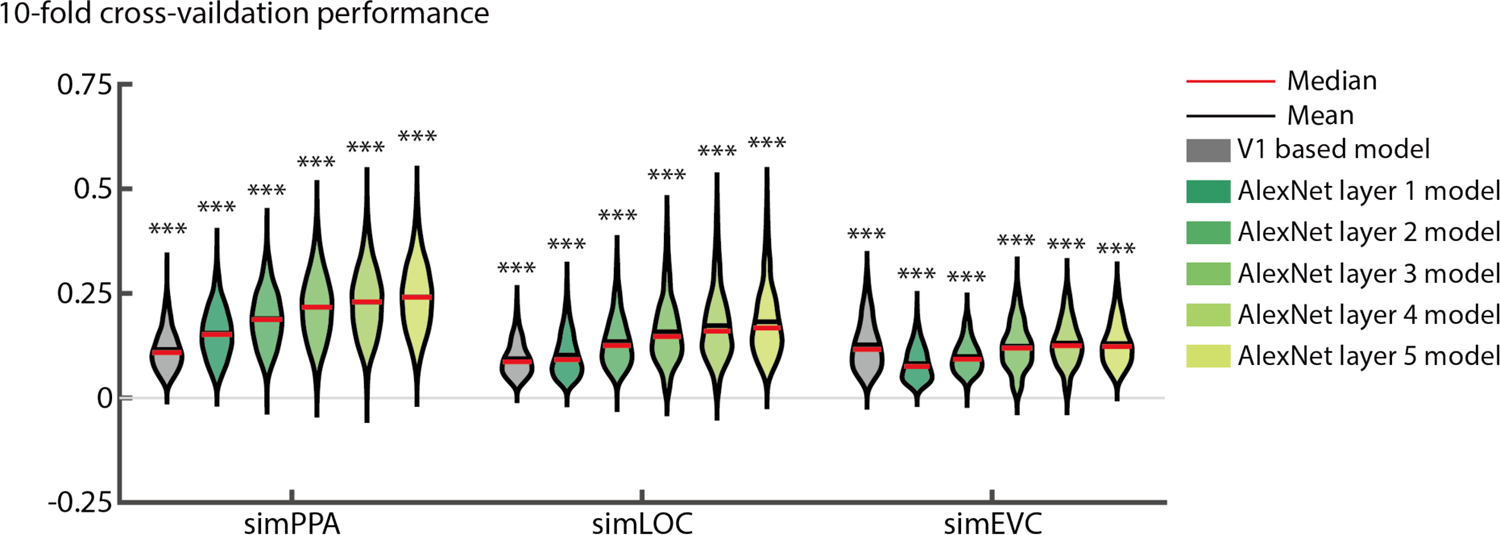
Performance of CNN encoding models. Voxelwise encoding models were trained using each convolution layer of AlexNet followed by global max pooling and LASSO regression. For comparison, encoding models were also constructed using a computational model of V1 instead of a CNN. Performance was assessed through 10-fold cross-validation on the BOLD5000 dataset. The average Pearson correlation of each voxel between the predicted and actual fMRI activations was computed across all folds of the cross-validation procedure. These violin plots show the distribution of encoding model performance across all voxels in all subjects for each ROI. ***p<0.001. All p-values were calculated by permutation tests (N=10,000 iterations). CNN: Convolutional neural network.

**Supplementary Figure 3.**
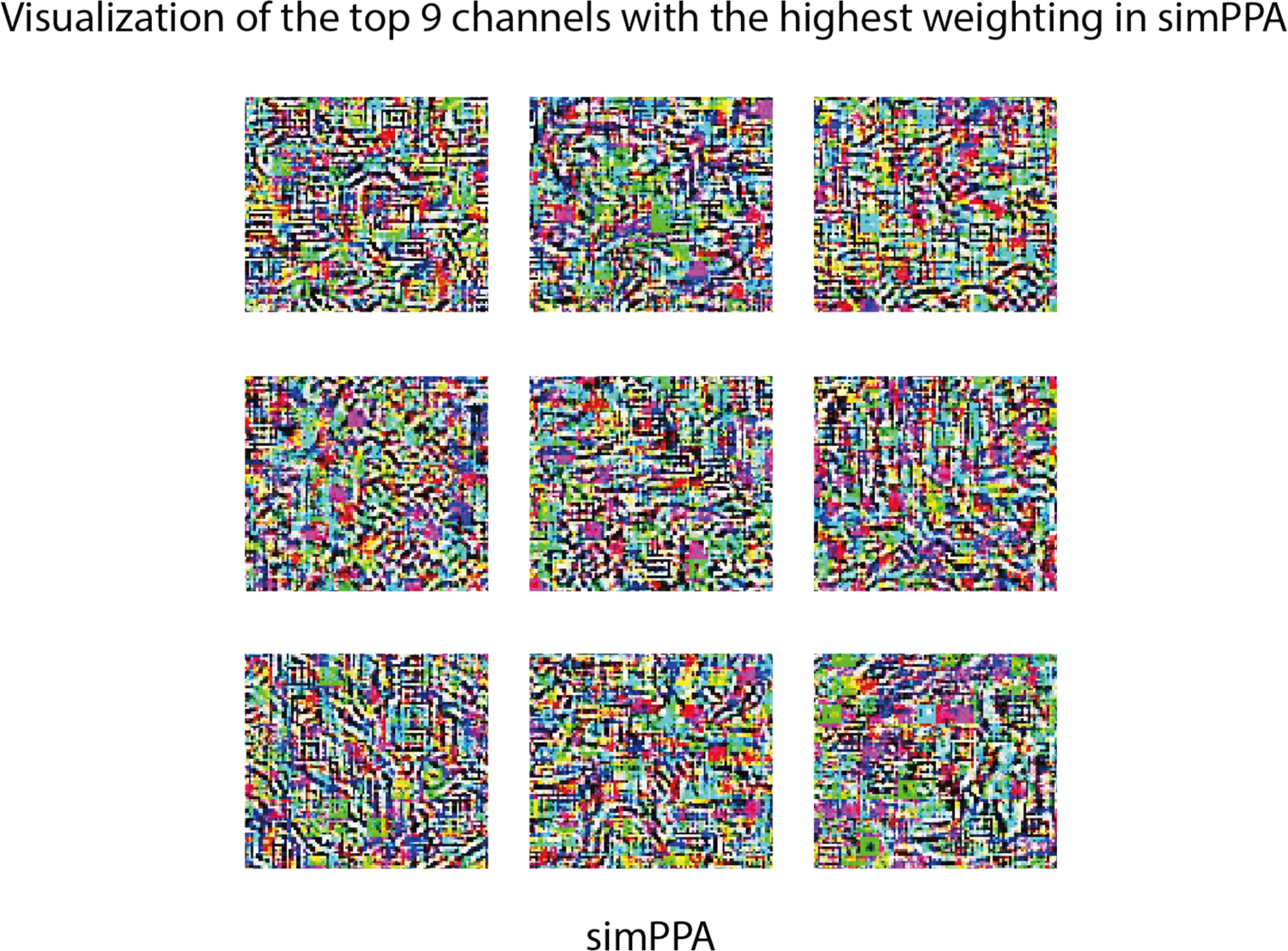
Visualization of CNN units with high encoding-model weights in simPPA. The top 9 channels from AlexNet layer 5 with the highest weightings in simPPA were visualized by synthesizing images that strongly drive the activation of these units. These units appear to encode complex visual features that cannot be easily described in words.

**Supplementary Figure 4.**
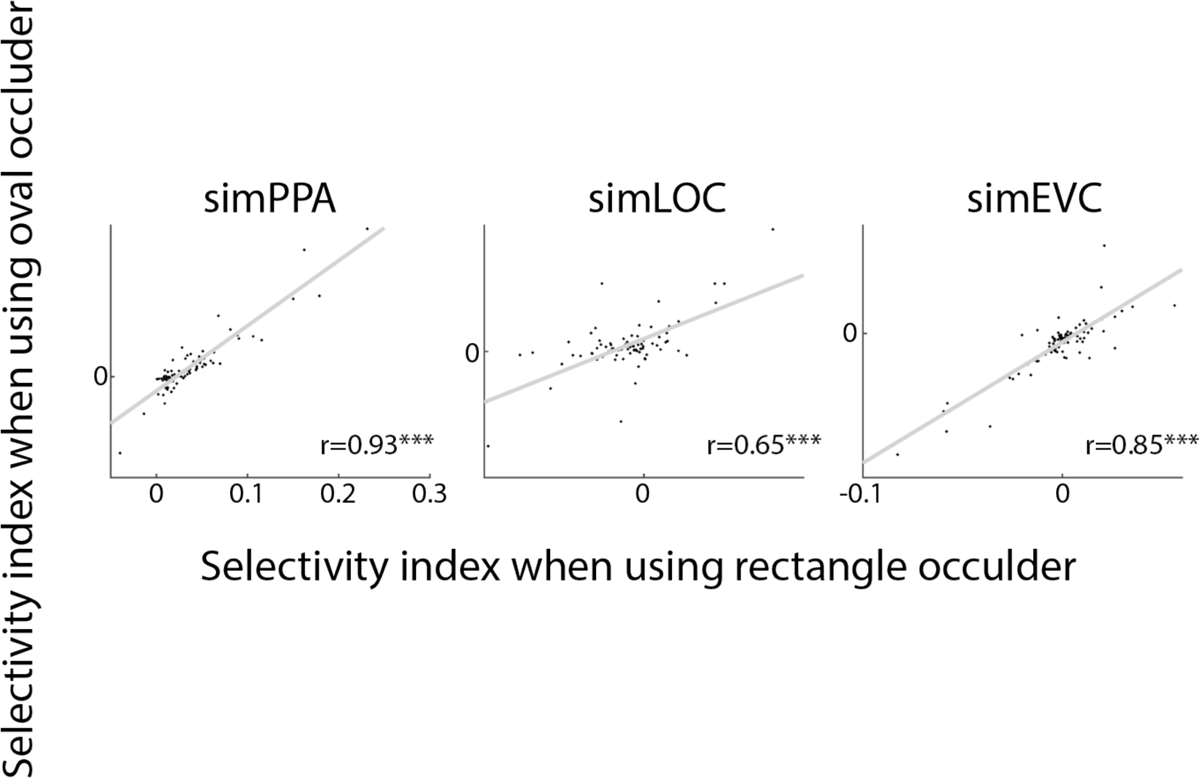
Robustness of the semantic preference mapping results to variation in occluder shape. Semantic preference mapping was conducted using both oval and rectangular occluders. These scatter plots show that for each simROI, the average selectivity indices from semantic preference mapping were highly similar regardless of whether the occluders were ovals or rectangles. ***p<0.001. All p-values were calculated by permutation tests (N=10,000 iterations).

**Supplementary Figure 5.**
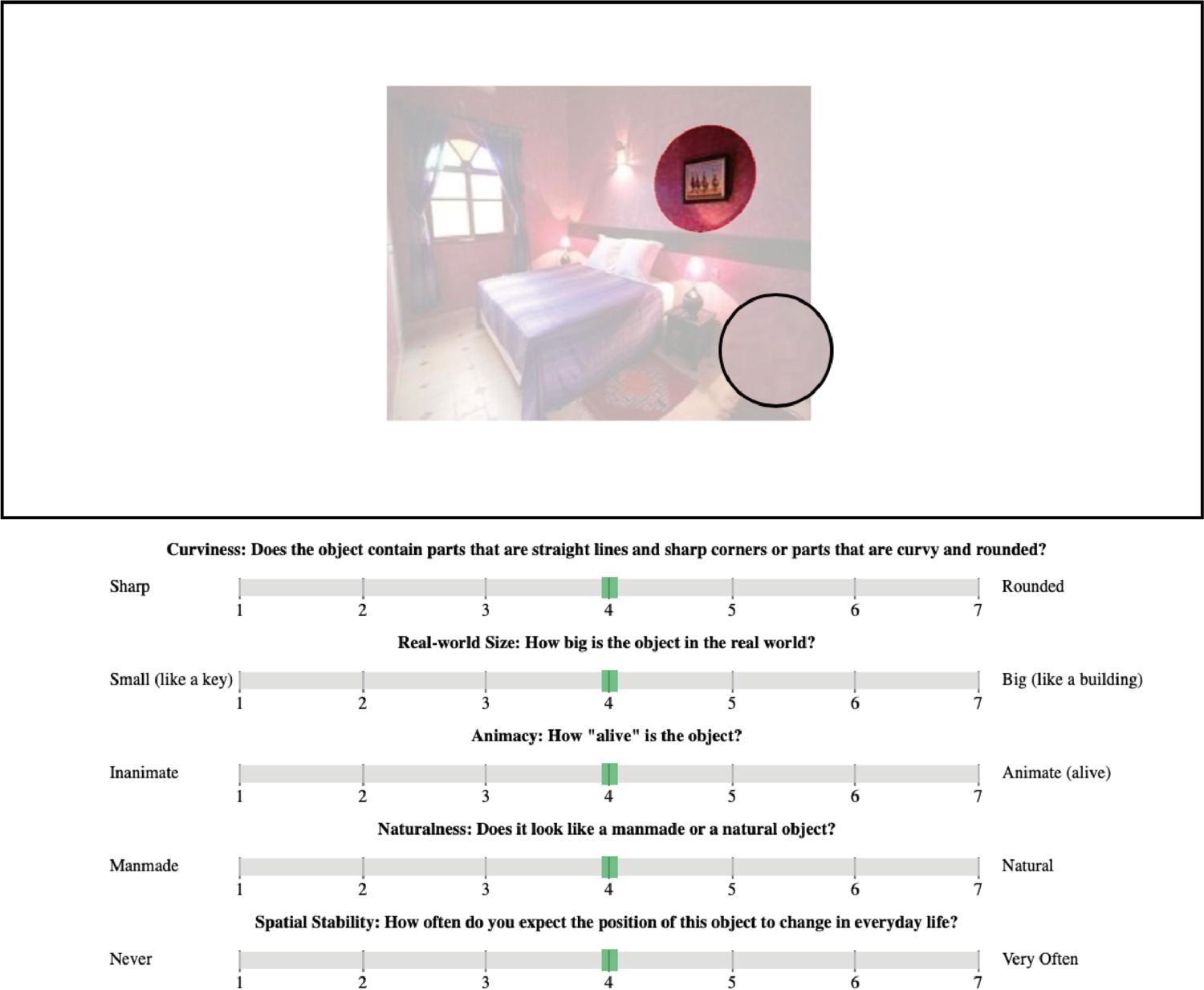
Object property ratings. The image depicts the webpage interface from the object property rating experiments. The target object was highlighted with a red oval, and the rest of the image was faded. A virtual magnifying glass could be moved around to enlarge portions of the image. Subjects were asked to provide ratings using a slider for five properties of the highlighted object.

**Supplementary Figure 6.**
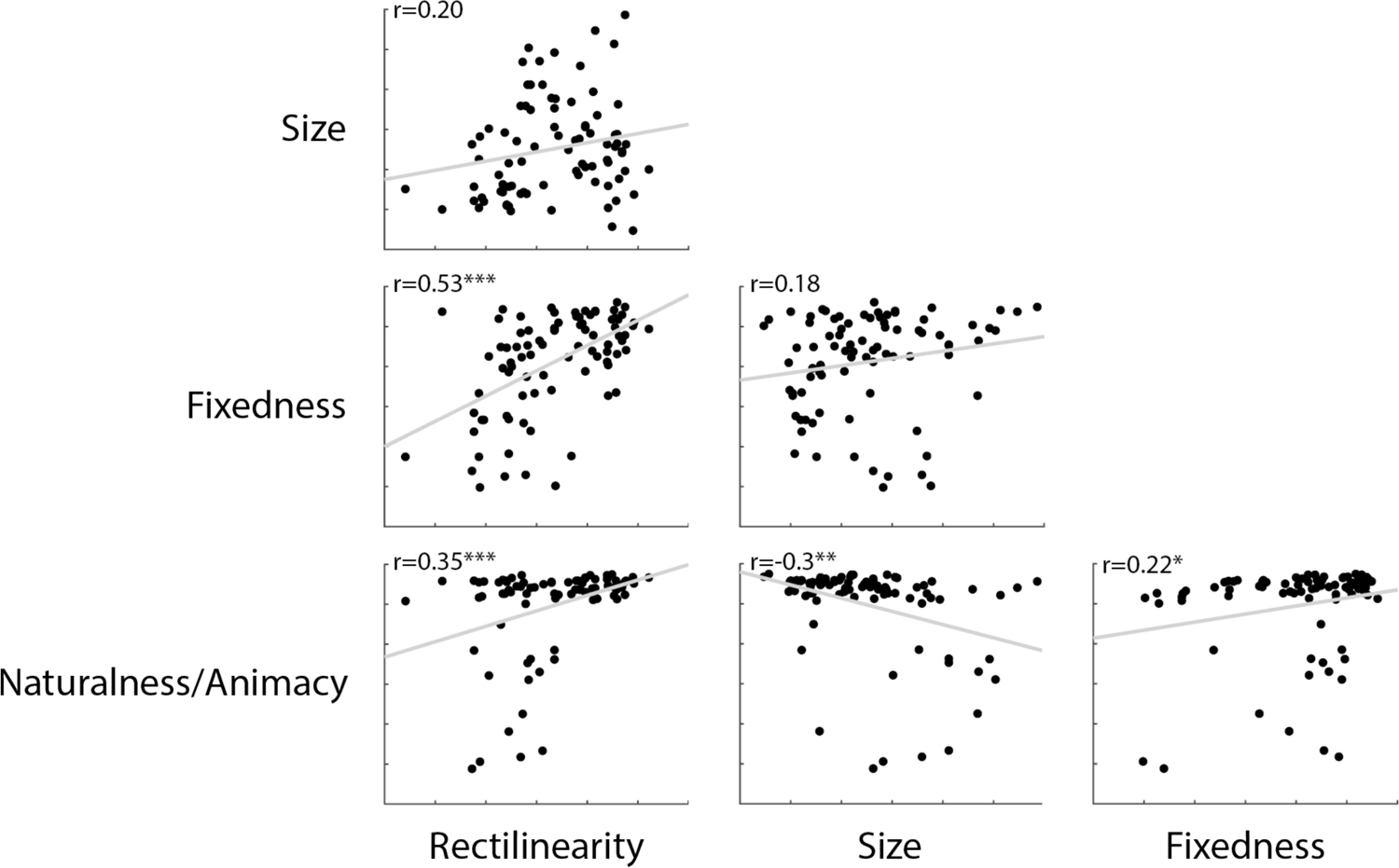
Covariance of object property ratings. These scatter plot show all pairwise correlations between the object properties. *p<0.05, **p<0.01, ***p<0.001. All p-values were calculated by permutation tests (N=10,000 iterations).

**Supplementary Figure 7.**
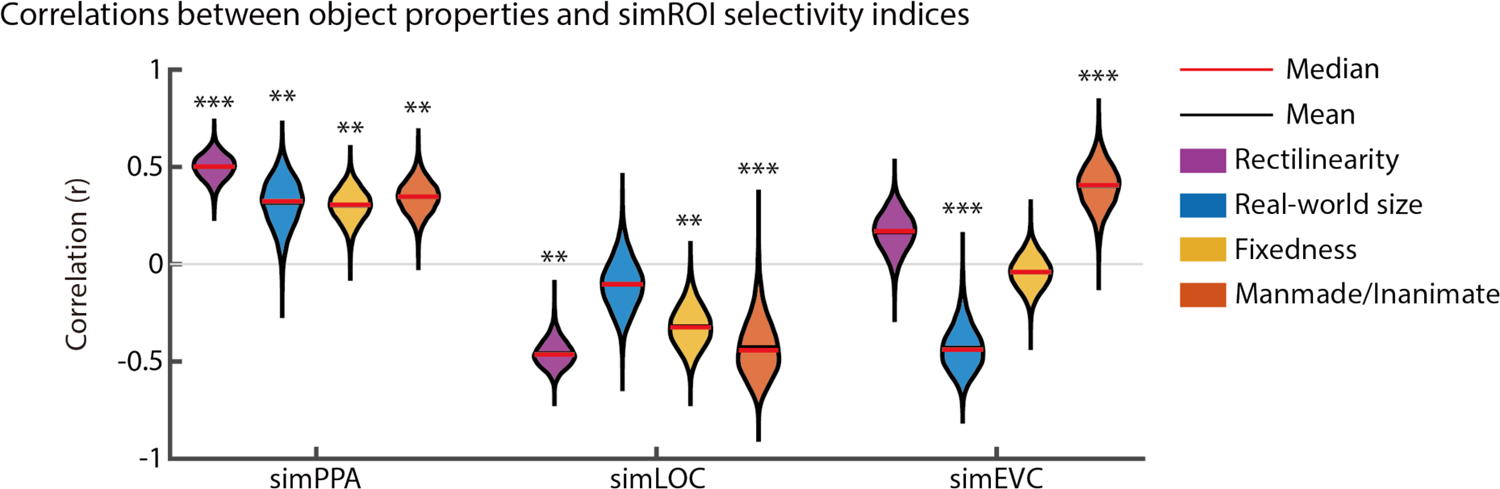
Correlations between object properties and selectivity indices for encoding models of different ROIs. This plot shows correlations between object properties and the average selectivity indices from multiple simROIs. All encoding models were created using layer 5 of AlexNet pre-trained on Places (as in *Fig. 4*). These results show that different simROIs exhibit different patterns of selectivity, even when they are built using the same CNN layer. The violin plots show distributions of the correlation values across 10,000 bootstrap resampling iterations. **p<0.01, ***p<0.001. All p-values were calculated by permutation tests (N=10,000 iterations).

**Supplementary Figure 8.**
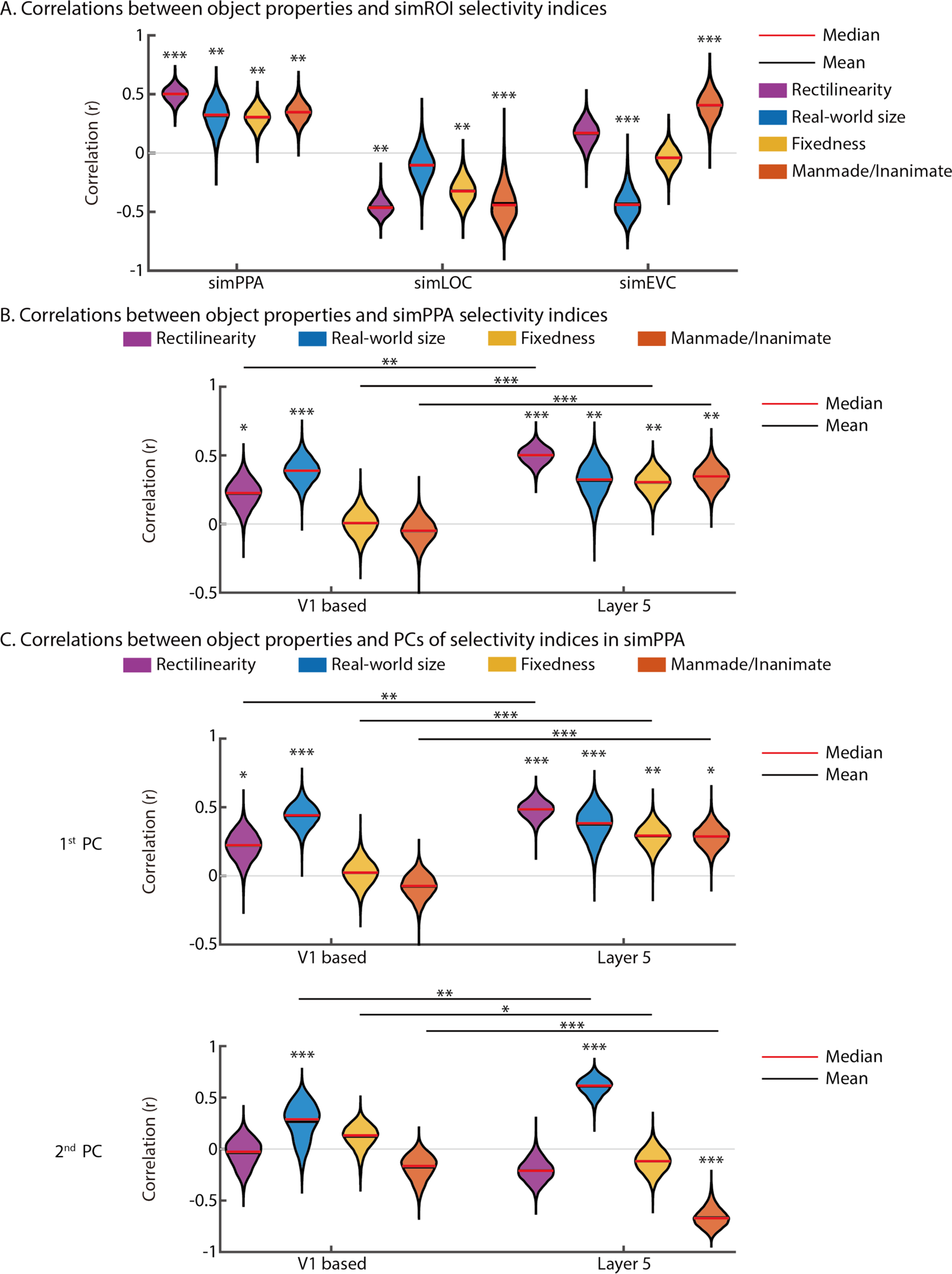
Comparing correlations between object properties and selectivity indices when building encoding models of the PPA from a CNN or a V1-like backbone. A) This plot shows correlations between object properties and the average selectivity indices from encoding models of the PPA using a CNN or a V1-like backbone. All correlations except for real-world size were significantly stronger in the CNN-based model. B) Similar findings held when examining the principal components of selectivity indices, with stronger correlations in the CNN-based model for all properties besides real-world size. These violin plots show distributions of the correlation values across 10,000 bootstrap resampling iterations. *p<0.05, **p<0.01, ***p<0.001. All p-values were calculated by permutation tests (N=10,000 iterations). PC: Principal component.

**Supplementary Figure 9.**
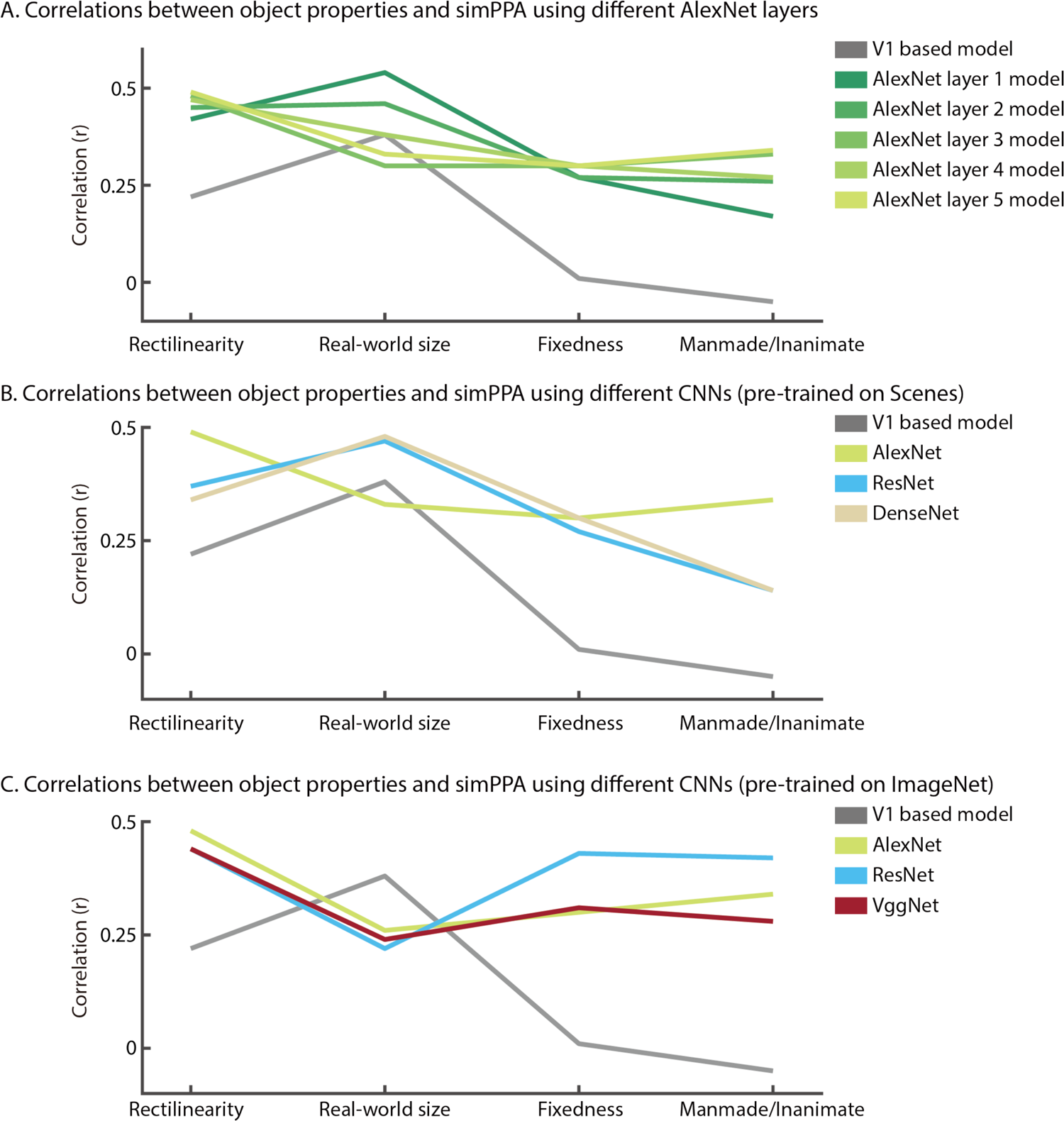
Visual feature tuning in simPPA gives rise to preferential responses to multiple high-level object properties when using different CNN backbones. These line plots show the mean correlations between object properties and selectivity indices for encoding models of the PPA built with different computational backbones. For each CNN, we examined the layer that produced the highest cross-validated encoding model accuracy in BOLD5000, and we verified that all encoding models exhibited significant generalization performance on the fMRI data from Bonner & Epstein, 2021 (all r>0.4 and all p<1e-4). DenseNet pre-trained on ImageNet failed the generalization test (r=0.03, p=0.79) and was thus not included in these analyses. A) PPA encoding models trained from different layers of AlexNet showed strong correlations with all four object properties. In comparison, all properties besides real-world size had weaker correlations in PPA encoding models built with a V1-like backbone. B) PPA encoding models built with different CNN architectures showed strong correlations with all four object properties. In comparison, all properties besides real-world size had weaker correlations in PPA encoding models built with a V1-like backbone. C) PPA encoding models built with CNNs pre-trained on the object images in ImageNet (instead of the scene images in Places) also showed strong correlations with all four object properties. In comparison, all properties besides real-world size had weaker correlations in PPA encoding models built with a V1-like backbone. CNN: convolutional neural network.

**Supplementary Figure 10.**
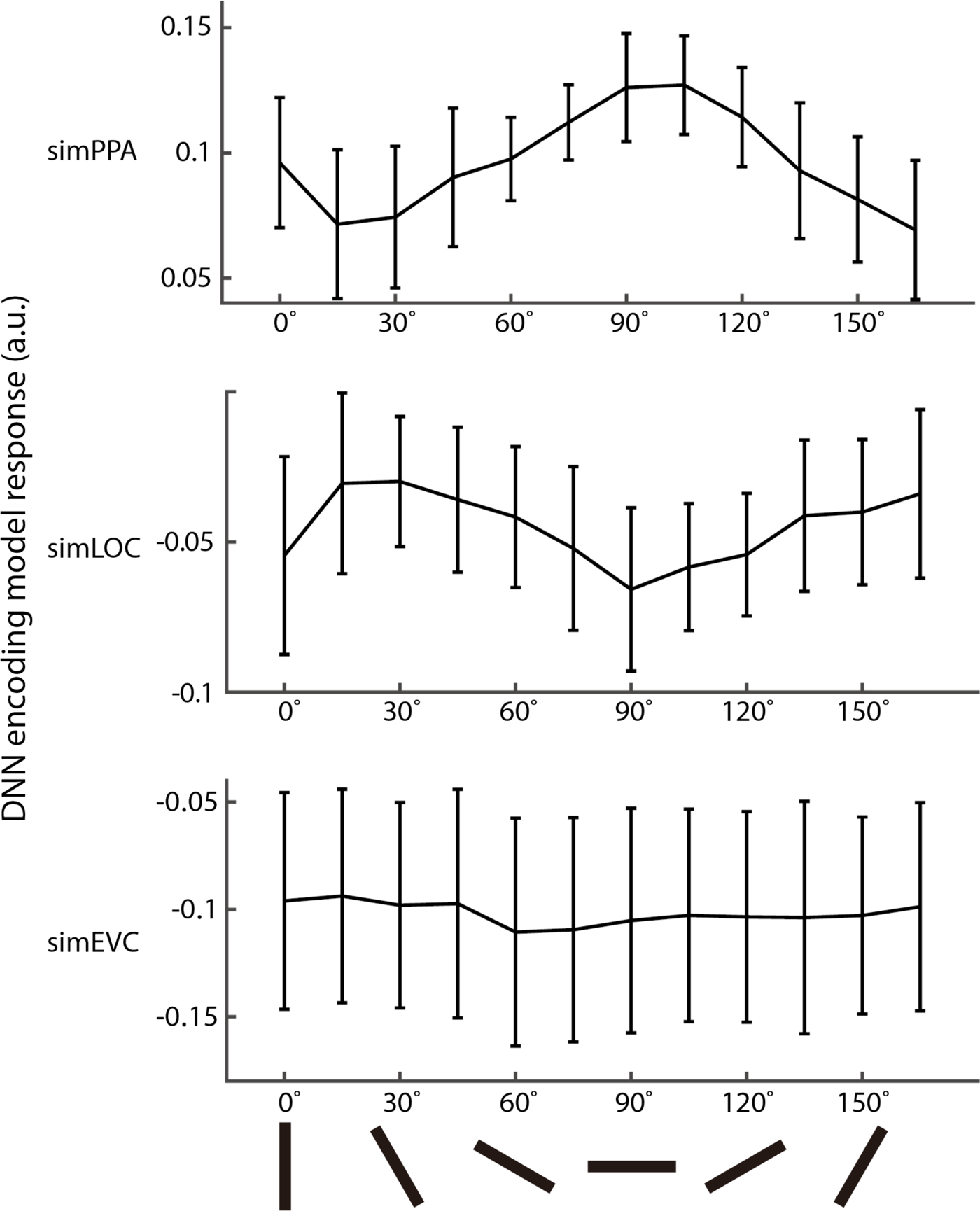
Encoding model responses to contour orientations. The average univariate response of each simROI is plotted for stimuli containing Gabor patches at a range of angles from 0° to 165°. Error bars represent +/-1 SD across the units of each simROI.

**Supplementary Figure 11.**
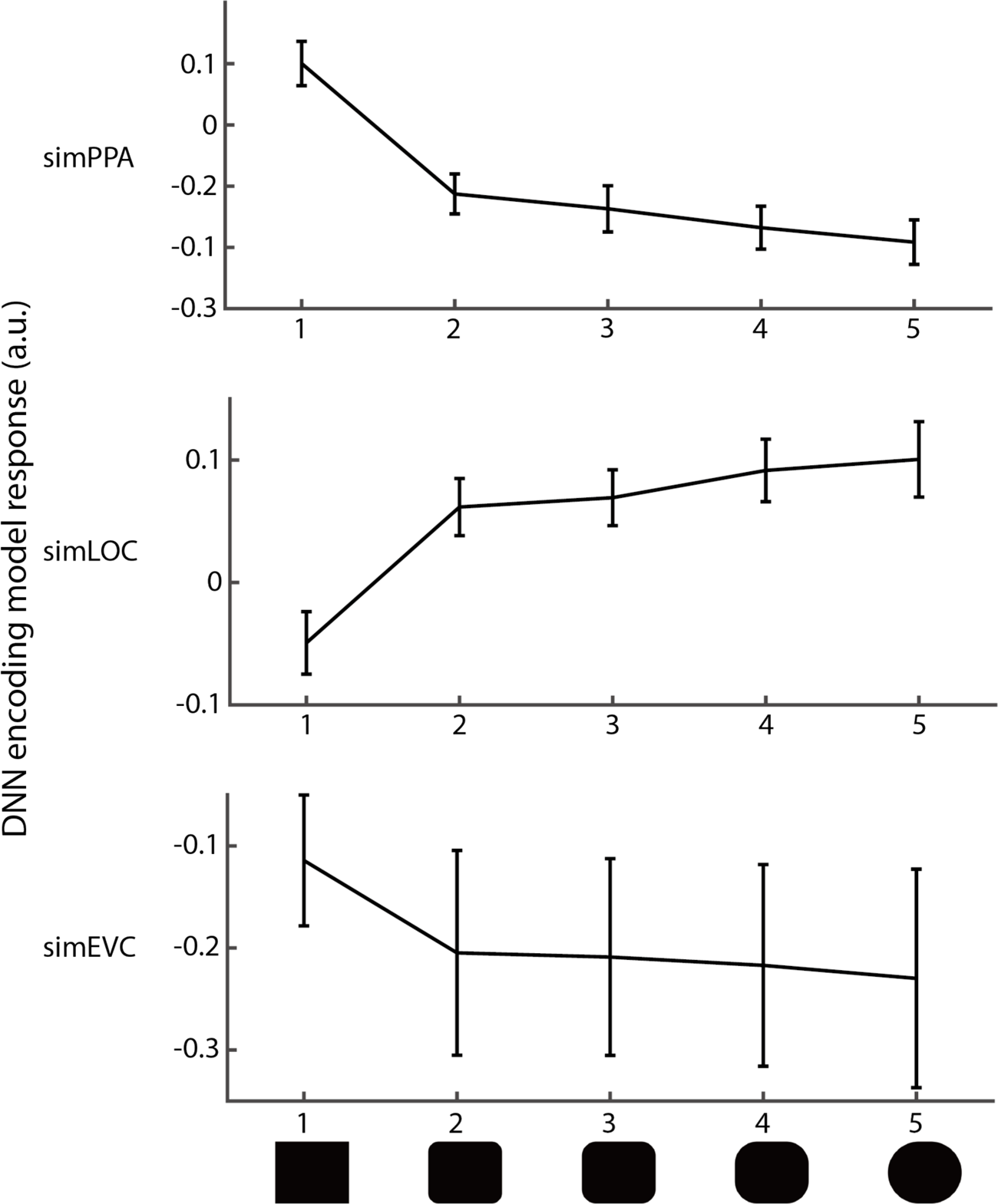
Encoding model responses to simple shapes. The average univariate response of each simROI is plotted for stimuli containing simple shapes that varied along a continuum from boxy to curvy. Error bars represent +/-1 SD across the units of each simROI.

**Supplementary Table 1.**
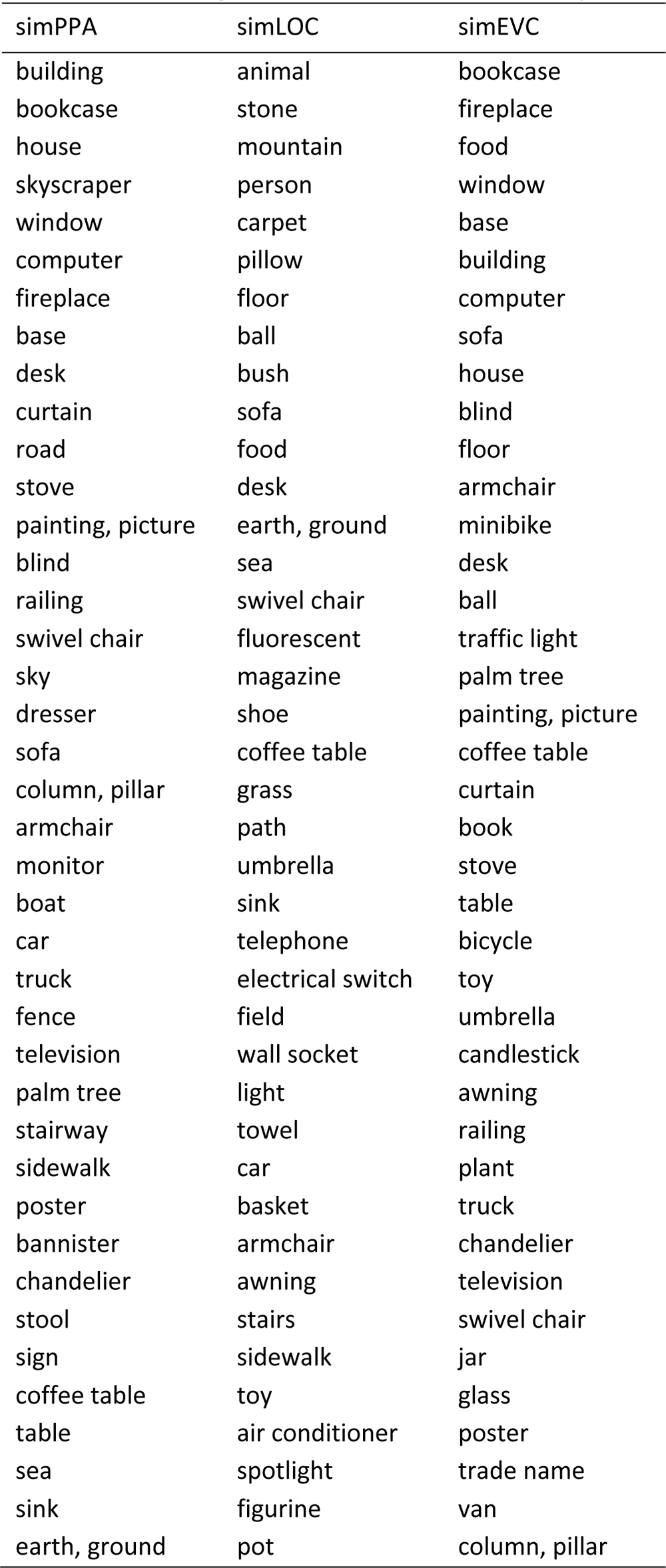
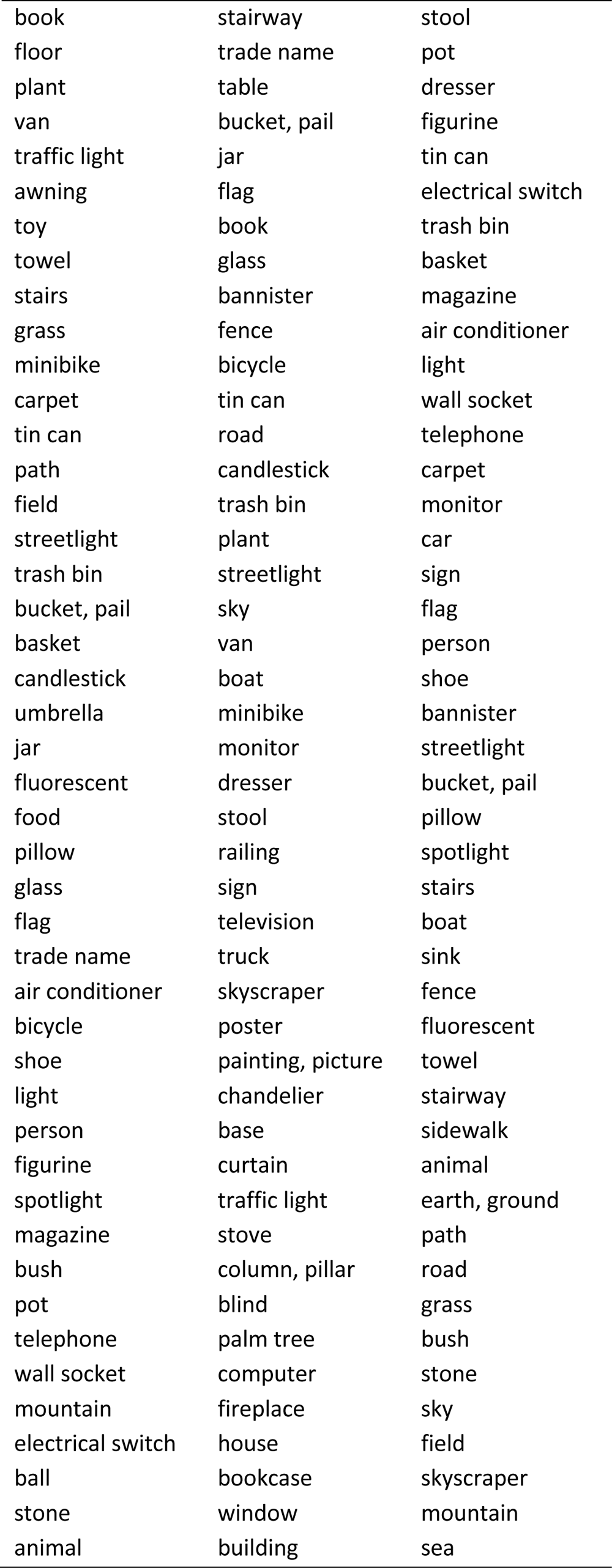
**Object categories used for semantic preference mapping.** This table shows all 85 object categories used in the semantic preference mapping procedure, with the objects sorted in descending order based on the selectivity indices for each simROI.

**Supplementary Table 2.** **simPPA trained on different neural networks.** Model performance and semantic occlusion results from simPPA trained with different layers from different neural networks. CV: 10-fold cross validation performance. SI: selectivity index. Rect.: rectilinearity. RWS: real-world size. Fix.: fixedness. Man.: manmade/inanimate.

|  | Performance | Correlation with SI |  |  |  |
| --- | --- | --- | --- | --- | --- |
| Correlation (r) | CV | Rect. | RWS | Fix. | Man. |
| AlexNet |  |  |  |  |  |
| L1 | 0.16 | 0.42 | 0.54 | 0.27 | 0.17 |
| L2 | 0.19 | 0.45 | 0.46 | 0.27 | 0.26 |
| L3 | 0.22 | 0.48 | 0.30 | 0.30 | 0.33 |
| L4 | 0.23 | 0.47 | 0.38 | 0.30 | 0.27 |
| L5 | 0.24 | 0.49 | 0.33 | 0.30 | 0.34 |
| ResNet |  |  |  |  |  |
| L1 | 0.17 | 0.42 | 0.46 | 0.25 | 0.21 |
| L2 | 0.21 | 0.43 | 0.30 | 0.28 | 0.24 |
| L3 | 0.25 | 0.45 | 0.41 | 0.34 | 0.25 |
| L4 | 0.26 | 0.37 | 0.47 | 0.27 | 0.14 |
| DenseNet |  |  |  |  |  |
| L1 | 0.22 | 0.43 | 0.46 | 0.31 | 0.20 |
| L2 | 0.24 | 0.48 | 0.34 | 0.30 | 0.29 |
| L3 | 0.26 | 0.38 | 0.46 | 0.31 | 0.18 |
| L4 | 0.26 | 0.34 | 0.48 | 0.30 | 0.14 |
| V1 model | 0.12 | 0.22 | 0.38 | 0.01 | -0.05 |

